# Pitfalls in performing genome-wide association studies on ratio traits

**DOI:** 10.1101/2023.10.27.564385

**Authors:** Zachary R McCaw, Rounak Dey, Hari Somineni, David Amar, Sumit Mukherjee, Kaitlin Sandor, insitro Research Team, Theofanis Karaletsos, Daphne Koller, Hugues Aschard, George Davey Smith, Daniel MacArthur, Colm O’Dushlaine, Thomas W Soare

**Affiliations:** Insitro, South San Francisco, CA, USA; Institut Pasteur, Université Paris Cité, Department of Computational Biology, Paris, FR; MRC Integrative Epidemiology Unit, University of Bristol, Bristol, UK; Centre for Population Genomics, Garvan Institute of Medical Research, Sydney, New South Wales, AU; Centre for Population Genomics, Murdoch Children’s Research Institute, Melbourne, Victoria, AU

**Author notes:** Correspondence: ZRM, TWS.

**Keywords:** Genome-wide association study, Polygenic scores, Proportion, Ratio, Body mass index, Collider bias

## Abstract

Genome-wide association studies (GWAS) are often performed on ratios composed of a numerator trait divided by a denominator trait. Examples include body mass index (BMI) and the waist-to-hip ratio, among many others. Explicitly or implicitly, the goal of forming the ratio is typically to adjust for an association between the numerator and denominator. While forming ratios may be clinically expedient, there are several important issues with performing GWAS on ratios. Forming a ratio does not “adjust” for the denominator in the sense of conditioning on it, and it is unclear whether associations with ratios are attributable to the numerator, the denominator, or both. Here we demonstrate that associations arising in ratio GWAS can be entirely denominator-driven, implying that at least some associations uncovered by ratio GWAS may be due solely to a putative adjustment variable. In a survey of 10 common ratio traits, we find that the ratio model disagrees with the adjusted model (performing GWAS on the numerator while conditioning on the denominator) at around 1/3 of loci. Using BMI as an example, we show that variants detected by only the ratio model are more strongly associated with the denominator (height), while variants detected by only the adjusted model are more strongly associated with the numerator (weight). Although the adjusted model provides effect sizes with a clearer interpretation, it is susceptible to collider bias. We propose and validate a simple method of correcting for the genetic component of collider bias via leave-one-chromosome-out polygenic scoring.

## Introduction

Ratio traits, those composed of a numerator trait divided by a denominator trait, are widely used in clinical practice, where they provide convenient scalar indicators of health and disease [1]. Examples include body mass index (BMI) in obesity [2]; forced expiratory volume (FEV1) to forced vital capacity (FVC) in chronic obstructive pulmonary disease (COPD) [3] and asthma [4]; phenylalanine to tyrosine concentrations in phenylalanine hydroxylase deficiency [5]; left ventricular ejection fraction (LVEF) in heart failure [6] and dilated cardiomyopathy [7]; serum aspartate (AST) to alanine aminotransferase (ALT) in hepatic cirrhosis [8]; and vertical cup-to-disc ratio (VCDR) in glaucoma [9]. As biomarkers of health, ratio traits have been used as targets for genome-wide association studies (GWAS) since the first applications of the method (e.g. BMI [10], serum metabolite ratios [11], WHR [12]), a practice that continues to the present (e.g. adipose tissue volumes [13], AST-ALT ratio [14], blood cell proportions [15], BMI [16], LVEF [17], lipid ratios [18], metabolite ratios [19, 20], protein ratios [21], seated to standing height [22, 23], skeletal proportions [24], urinary albumin-creatinine ratio (UACR) [25], VCDR [26, 27], WHR [28]). While simple ratios may provide convenient heuristics for making patient care decisions, this does not imply that ratios are the ideal, or even an appropriate, vehicle for understanding complex trait genetics.

The use of ratio variables in regression models has several well-known statistical issues [29]. Explicitly or implicitly, the goal of forming a ratio is generally to control for an association between the numerator and denominator. However, forming a ratio does not “adjust” for the denominator in the sense of conditioning on it. Moreover, the genetic effect estimated by the ratio model is difficult to interpret because the association may be due to the numerator, the denominator, or both. An alternative to the ratio model is to condition on the denominator as a covariate, which we refer to as the “adjusted model.” This model isolates the direct effect of genotype on the numerator while holding the denominator constant. However, conditioning on a heritable covariate introduces the possibility of collider bias [30, 31].

Here we demonstrate several pitfalls that can arise when performing GWAS of ratio traits.

As a running example, we consider BMI, the ratio of weight in kilograms to height^2^ in meters^2^ (**Figure S1**). Through simulations and analyses of real data in which the numerator (i.e. weight) is permuted, we demonstrate that associations with ratio traits can be entirely denominator-driven, and that the probability of detecting such associations increases with denominator heritability. Thus, at least some associations with BMI are likely attributable to height. In a survey of 10 common ratio traits, we find that the ratio and adjusted models disagree at roughly 1/3 of loci, and that associations detected by only the ratio model are enriched for denominator-driven signals. Variants identified only by the ratio model tend to be more strongly associated with height (the denominator), while variants identified only by the adjusted model tend to be more strongly associated with weight (the numerator). We scrutinize the practice of correlating effect sizes from two different models (e.g. ratio and adjusted) fit to the same data set, and show that doing so tends to overstate the agreement of those models. Finally, we address the issue of heritable covariate bias and propose a method of correcting for the component of collider bias that arises from common genetic causes.

## Results

### Ratio model associations can be entirely denominator-driven

Figure 1(a) presents a simple schematic of how a genetic variant can influence a ratio via an effect on the denominator only. To empirically demonstrate that associations obtained from ratio GWAS can be entirely denominator-driven, we conducted analyses of a null phenotype, permuted body weight, within the the UK Biobank (UKB; *N* = 356K) [32, 33]. Permutation keeps the mean and standard deviation (SD) of weight unchanged but abolishes any association with genotype or the denominator. Figure 1(c) compares Manhattan plots for GWAS of weight (unpermuted), height, permuted weight adjusted for height and height^2^ (the “adjusted model”), and the ratio of permuted weight to height^2^ (the “ratio model”). As desired, the adjusted model detects no genome-wide significant (GWS) loci (*P ≤* 5 *×* 10^−8^) [34]. By contrast, the ratio model detects 142 independent GWS loci, and exhibits clear similarity to the marginal Manhattan plots for height. Of these 142 GWS loci, 140 (98.6%) were GWS for height, and the remaining 2 were suggestively significant (*P ≤* 1 *×* 10^−6^). Transformation of the ratio, via the logarithm or the rank-based inverse normal transform [35], does not prevent the detection of denominator-driven associations (**Figure S2**). Because the numerator was permuted, and consequently had no association with genotype, the signals detected by the ratio model in this experiment must be entirely attributable to the denominator.

**Figure 1:**
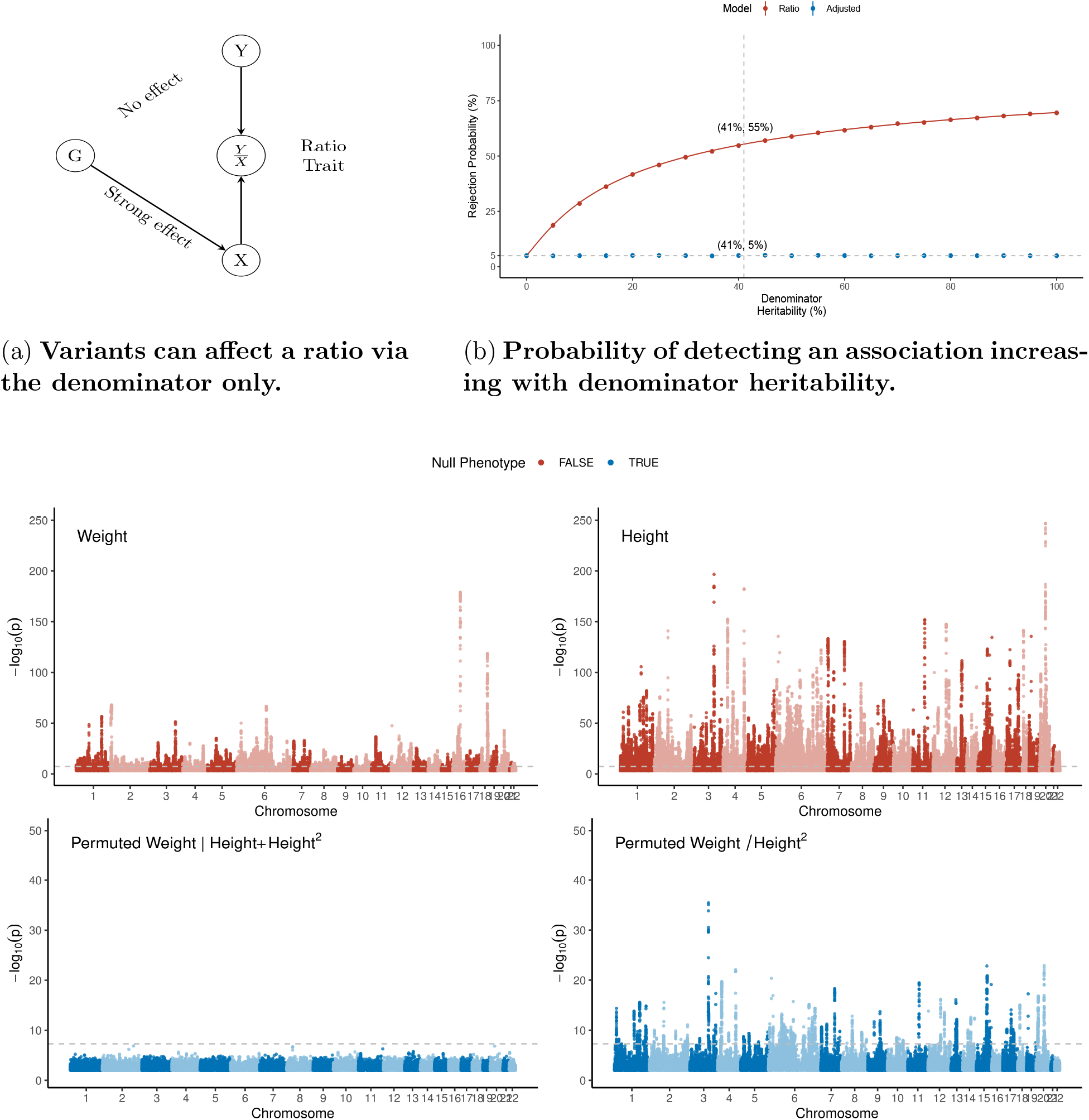
Simulated and empirical evidence that associations detected in ratio GWAS can be entirely denominator-driven. (a) Example data generating process in which a variant *G* affects a ratio phenotype *Y/X* by influencing the denominator *X* only. (b) Simulation study (*N* = 10K) of rejection probability as a function of denominator heritability. The numerator was permuted weight, and the denominator was simulated to control the heritability while matching the mean and variance to the observed values for height^2^. Each dot is represents the mean of 10^5^ simulations. (c) Manhattan plots for GWAS conducted in the UK Biobank (*N* = 356K). Each dot represents a genetic variant.

### Propensity to detect denominator-driven associations increases with denominator heritability

To study the effect of denominator heritability on the probability of observing an association with a ratio phenotype, a random sample of 10K subjects was drawn from the UKB. The numerator was permuted body weight. To enable control of the heritability, the denominator was simulated from an infinitesimal model in such a way that the mean and SD matched the observed values for height^2^ in UKB (Supplemental Methods). Figure 1(b) tracks the rejection probability for the ratio and adjusted models. The probability that the adjusted model detects an association remains flat at the type I error rate of 5%, which is expected given that the numerator is unassociated with genotype. By contrast, the probability that the ratio model detects an association grows in concert with the heritability of the denominator (representing height^2^). The solid red line is a theoretical calculation of the expected rejection probability as a function of denominator *h*^2^ (Supplemental Methods). At the empirical heritability for height, estimated by linkage-disequilibrium (LD) score regression [36] in the full cohort, the ratio model detected an association with 55% probability. This experiment demonstrates that the risk of detecting denominator-driven associations increases with the denominator heritability. The interpretation of associations detected by the ratio model is further complicated by the fact that the number of association detected depends on the mean of the numerator, even when the numerator is null (**Figure S3**). Additional simulations reported in Supplemental Sections 2.5 and 2.6 examine the effects of environmental correlation and pleiotropy on the operating characteristics of the ratio and adjusted models.

### Ratio and adjusted models can reach discrepant conclusions

We surveyed the extent to which ratio traits are used in the literature by querying the NHGRI-EBI GWAS Catalog [37] for trait names including: concentration, fraction, index, percent(age), percentile, proportion, rate, and ratio; as well as specific traits known to be ratios (e.g. BMI, WHR). We manually reviewed the 667 traits identified for evidence of having a heritable denominator, excluding, for example, per-volume concentrations or per-time rates. Using this approach, we ascertained that 362 traits, representing 3.2% of all traits and 7.8% of all reported associations in the GWAS Catalog, involved ratios with a heritable denominator (Supplementary Data).

To understand the extent to which the ratio and adjusted models differ in real data, we conducted two GWAS within the UKB on each of 10 phenotypes analyzed as ratios: one with the ratio as the outcome and the other with the numerator as the outcome, adjusting for the denominator (the “adjusted” analysis). The numbers of genome-wide significant (GWS) loci identified by the ratio and adjusted models are presented in **Table 1**. The union of loci detected by the ratio and adjusted models was partitioned into 3 sets (ratio-only, adjusted-only, or both) based on whether or not the lead variant from each ratio locus was within 250kb of the lead variant for any adjusted locus, and vice versa. On average the ratio model detected 386 loci while the adjusted model detected 292 loci. That the ratio model tends to identify more loci is expected given that a variant may associate with the ratio either through the numerator or the denominator. On average, 67.6% of all loci detected were detected by both models, 22.8% were unique to the ratio model, and 9.6% were unique to the adjusted model. Thus, although the ratio and adjusted models are generally in agreement, for 32.6% of loci the two analyses may reach differing conclusions. We also assessed the percentage of signals from the ratio and adjusted models that were within 250kb of a lead variant for the denominator trait, but not within 250kb of a lead variant for the numerator trait (“% denominator-driven”). On average, 48% of ratio-only signals were shared with the denominator trait, while only 8% of adjusted-only signals were shared with the denominator trait. Thus loci detected by only the ratio model are enriched for denominator-driven associations.

**Table 1:** Survey of discrepancies between the ratio and adjusted models across traits analyzed as ratios. Number of independent genome-wide significant loci detected by the ratio and adjusted models, categorized by whether the signal was detected by the ratio model only or the adjusted model only. Loci genome-wide significant for the denominator, but not the numerator, were considered “denominator-driven.” Loci were obtained by clumping full-genome summary statistics at *R*^2^ *≤* 0.10 and *P ≤* 5 *×* 10^−8^.

| Traits |  |  | GWS Loci |  |  |  |
| --- | --- | --- | --- | --- | --- | --- |
| Numerator | Denominator | $N$ | Ratio | Adjusted | Ratio-only<br>(% Denominator-driven) | Adjusted-only<br>(% Denominator-driven) |
| Albumin (blood) | Creatinine (blood) | 311,122 | 944 | 556 | 727 (71%) | 314 (0%) |
| Albumin (urine) | Creatinine (urine) | 105,501 | 6 | 4 | 6 (0%) | 4 (0%) |
| Weight | Height<sup>2</sup> | 355,627 | 844 | 830 | 89 (34%) | 70 (4%) |
| FEV<sub>1</sub> | FVC | 325,220 | 538 | 474 | 86 (13%) | 49 (2%) |
| Stroke volume | End diastolic volume | 29,171 | 6 | 0 | 6 (0%) | 0 (.) |
| Phenylalanine | Tyrosine | 200,159 | 251 | 174 | 54 (56%) | 10 (0%) |
| Platelets | Lymphocytes | 52,133 | 2 | 0 | 2 (0%) | 0 (.) |
| Seated height | Standing height | 355,373 | 440 | 115 | 330 (45%) | 39 (8%) |
| Trunk fat mass | Whole body fat mass | 349,533 | 359 | 240 | 132 (0%) | 46 (0%) |
| Waist circumference | Hip circumference | 356,112 | 465 | 529 | 116 (1%) | 115 (37%) |
| Average |  | 243,995 | 385.5 | 292.2 | 154.8 (48%) | 64.7 (8%) |

### Effect size correlation overstates concordance of ratio and adjusted models

Several studies of ratio phenotypes have performed sensitivity analyses in which the effect sizes from a ratio model are compared with those from an unadjusted model [10, 12, 17, 13] or an adjusted model [24]. Comparisons among effect sizes from different models fit to the same data should be interpreted with caution. As demonstrated analytically in the Supplemental Methods, such estimates are inherently correlated due to having been obtained from the same subjects. To enable an unbiased comparison of effect sizes across models, we split the full UKB cohort (*N* = 356K) into two equally sized subsets, A and B (*N* = 178K each). Subset B serves as the validation cohort for associations discovered in subset A and vice versa. Importantly, effect sizes estimated from models fit in A can fairly be correlated with effect sizes estimated in B.

Figure S5 presents the effect size correlations between BMI and a comparator GWAS where the effect sizes were either estimated in the same or in different (split) samples.

The comparators include marginal GWAS of height and weight, as well as GWAS of weight adjusted for height (and height^2^). Although correlation remains, the magnitude of correlation declines appreciably when the effect sizes are measured in independent samples. For example, in the case of weight adjusted for height, the correlation with BMI drops from 97.8% to 81.1%.

### Ratio associations are enriched for denominator signal

Comparing the ratio and adjusted models enables variants to be partitioned into 3 classes: variants detected by the ratio model only (“ratio-only”), variants detected by the adjusted model only (“adjusted-only”), and variants detected by both (“both”). To understand the biological significance of these three groups of variants, in **Figure S6** we explore the enrichment of signal from marginal associations, calculated in the full UKB cohort, for the numerator (weight) and denominator (height) traits at these variants. Consistent with the hypothesis that variants detected by the ratio model only are more likely to be denominator-driven (and that variants detected by the adjusted model only are more likely to be numerator-driven), the mean *χ*^2^ of height was larger at ratio-only loci (two-sided T-test, *T* = 3.21; *p* = 0.0015), whereas the mean *χ*^2^ for weight was larger at adjusted-only loci (*T* = 3.19; *p* = 0.0017).

We also compared the power of the ratio and adjusted models at GWS loci for obesity-related traits compiled from external sources (excluding the UKB cohort where possible): body fat percentage [38], chronically-elevated alanine aminotransferase (cALT) [39], coronary artery disease (CAD) [40], hip circumference [41], low-density lipoprotein (LDL) cholesterol [42]), and type 2 diabetes (T2D) [43]. The adjusted model had larger *χ*^2^ statistics at variants defined by several obesity-relevant traits, including body fat percentage (paired two-sided T-test, *T* = 2.42; *p* = 0.033), hip circumference (*T* = 8.4; *p* = 4.3 *×* 10^−13^), and LDL cholesterol (*T* = 2.67; *p* = 0.0074, **Table S4**). Conversely, in no case did the ratio model have larger *χ*^2^ statistics. Together, these observations suggest that GWAS of weight adjusted for height may provide a more direct and powerful route to understanding the genetic basis of adverse adiposity.

### Leave-one-chromosome-out polygenic scoring can partially correct for heritable covariate bias

The denominators of the ratios considered in **Table 1** are themselves heritable (**Table S3**). Important work by [30, 31] underscores that adjusting for a heritable covariate can introduce collider bias. Consider the data generating processes represented in Figure 3(a). Here *Y* is the phenotype, *G* is genotype, *X* is a covariate, and *B* represents background common causes of *X* and *Y*. Supposing *B* has an effect on both *X* (*α_B_* ≠ 0) and *Y* (*β_B_* ≠ 0), if *G* has an effect on *X* (*α_G_* ≠ 0), then conditioning on *X* opens a backdoor path [44] between *G* and *Y*, namely 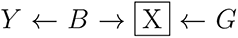. The presence of this backdoor path biases estimation of *β_G_*, the genetic effect of interest. In cases where *G* has no effect on *X* (*α_G_* = 0), *X* is no longer a collider and the bias vanishes. Supposing *α_G_* ≠ 0, another way to remove collider bias would be to condition on *B*. When *B* is genetic background, this can be achieved by performing conditional GWAS for *Y* adjusting for all variants that affect *X*. In practice, adjusting for all variants affecting *X* may be impractical. Instead, we pursue the strategy of conditioning on a polygenic score (PGS) for *X*, effectively blocking the path from *B → X*.

**Figure 3:**
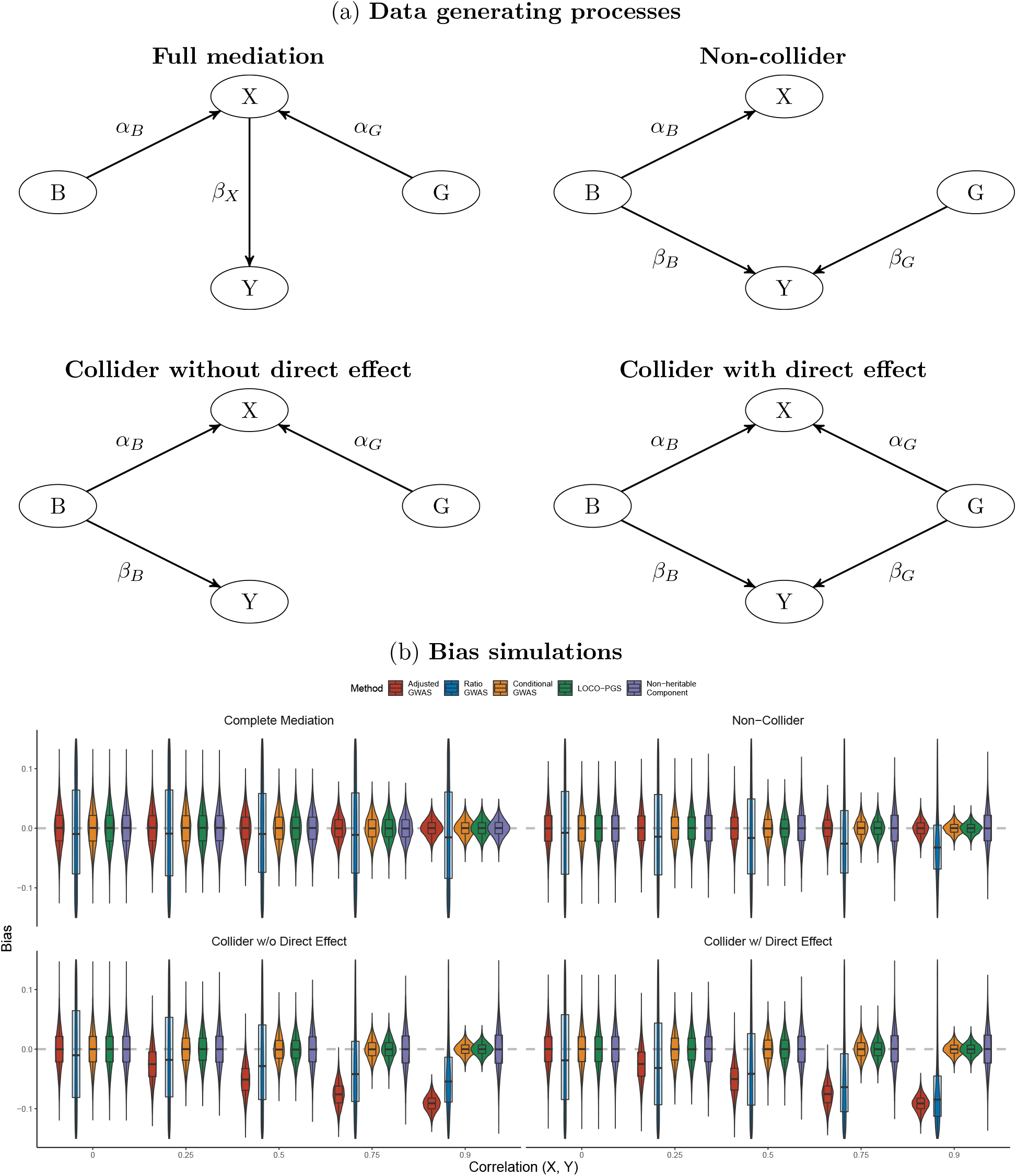
Leave-one-chromosome-out polygenic scoring corrects for collider bias due to the genetic background. (a) Data generating processes considered in the simulation study. In each case, *B* is genetic background, *G* is genotype, *X* is a heritable covariate, and *Y* is the phenotype of interest. (b) Bias in estimating *β_G_* as a function of the correlation between *X* and *Y* due to *B*. The sample size is *N* = 10^3^ and each violin presents the distribution of 10^4^ simulations replicates. The lower, middle, and upper bars of the box plots demarcate the 25th, 50th, and 75th percentiles, and the whisker extend to 1.5× the inter-quartile range. Adjusted and ratio are susceptible to bias; conditional is unbiased and theoretically optimal; LOCO-PGS and non-heritable component are the proposed bias-correction methods.

Drawing on ideas from BOLT-LMM [45] and Regenie [46], we propose constructing a leave-one-chromosome-out (LOCO) PGS for the heritable covariate *X*, then performing GWAS adjusting for both *X* and its LOCO-PGS. A related strategy is to regress the LOCO-PGS out of the heritable covariate then adjust for the residual “non-heritable” component (in practice, the residual component may retain some heritability if the PGS does not capture all genetic contributions to the heritable covariate [47]). In either case, the PGS serves to correct for collider bias by breaking the association between the genetic background and the heritable covariate. In the Supplemental Methods, we demonstrate analytically that excluding variants in LD with the focal variant is necessary (i.e. adjusting for a global rather than a LOCO PGS would introduce bias), and that LOCO-PGS adjustment successfully removes collider bias due to the genetic background. Aschard *et al* [30] demonstrated that the magnitude of heritable covariate bias is proportional to ℂ(*Y, X*) *·* ℂ(*X, G*), where ℂ(*·, ·*) denotes the covariance. In connection with their result, the LOCO PGS attenuates the component of ℂ(*Y, X*) that is due to genetic background. When the covariance between *Y* and *X* is solely the result of genetic factors, the LOCO PGS has the potential to fully eliminate the heritable covariate bias.

Figure 3(b) empirically assess the bias of adjusted, ratio, and conditional GWAS alongside the proposed methods of correcting for genetic background under each data generating process in Figure 3(a). Adjusted GWAS is unbiased when the heritable covariate *X* fully mediates the effects of genotype *G* and genetic background *B* on the phenotype, or when genotype has no effect on *X*, but incurs bias when *X* is a collider. Conditional GWAS is the optimal strategy of controlling collider bias, but requires conditioning on *B*. Both LOCO-PGS and non-heritable covariate adjustment remove bias due to the genetic background, but LOCO-PGS is more efficient (i.e. produces smaller standard errors). Ratio GWAS is often biased, and inefficient, because the association model is misspecified. Specifically, genotype affects the numerator and/or denominator individually, not as a ratio.

Figure 4 compares validation effect sizes from several models with discovery effect size from the adjusted model using real data from the UKB. The similarity of the effect sizes from the adjusted, LOCO-PGS, and non-heritable component models suggests either that most variants detected by the adjusted model are not subject to substantial collider bias, or that the collider bias is due primarily to factors other than genetic background, for which the LOCO-PGS cannot account.

**Figure 4:**
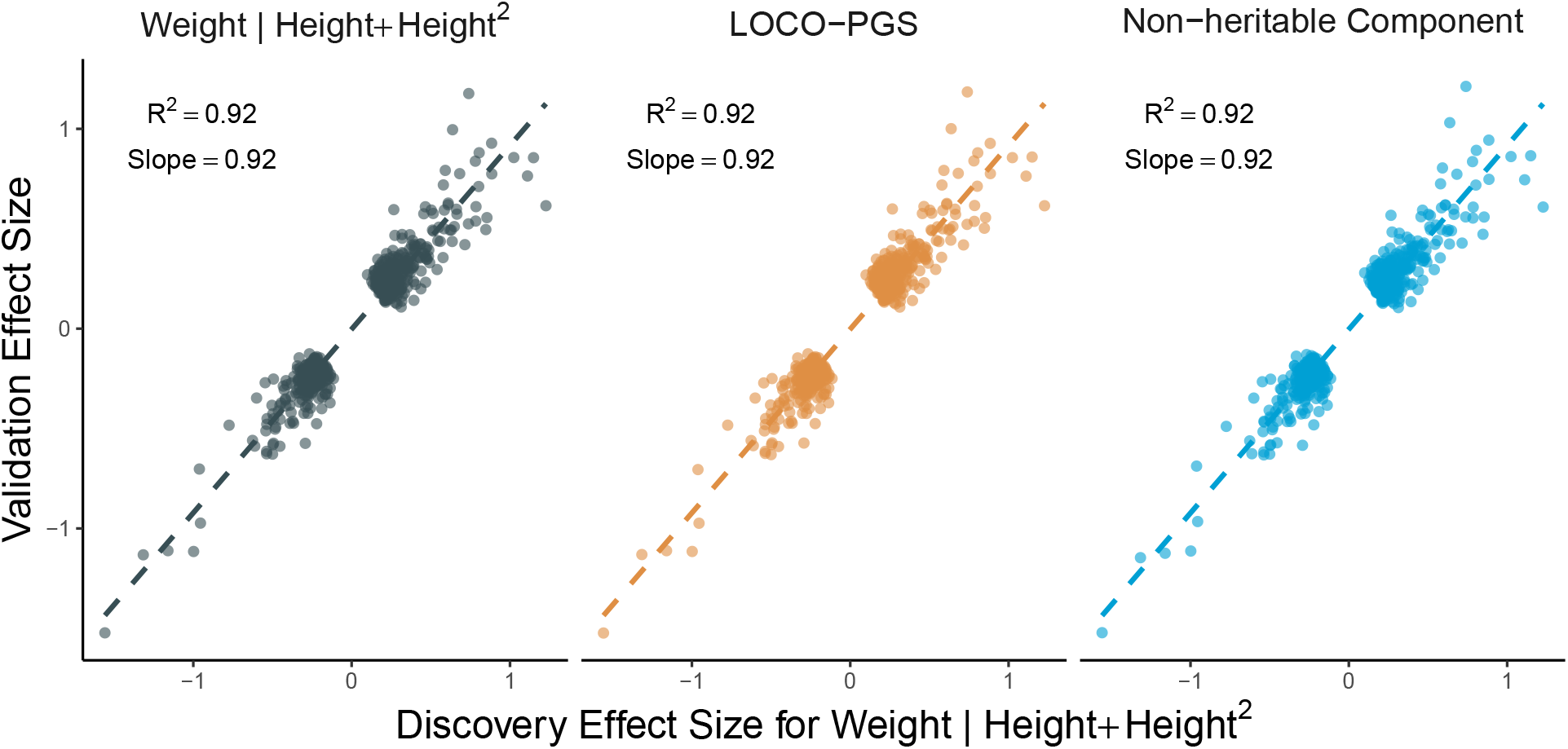
Empirically, effect sizes from the adjusted and LOCO-PGS analyses are similar. Validation effect sizes from several GWAS are compared with the discovery effect size from the adjusted model, at independent (*R*^2^ ≤ 0.1) genome-wide significant (*P* 5 10^−8^) loci for the latter. The discovery and validation effect sizes were estimated in two independent subsets of the UK Biobank (*N* = 178K each). The adjusted model performs GWAS of weight adjusting for height and height^2^. The LOCO-PGS model performs GWAS of weight adjusting for height and a leave-one-chromosome-out polygenic score (LOCO-PGS) for height. The non-heritable component model performs GWAS of weight adjusting for the residual after regression height on its LOCO-PGS.

## Discussion

Ratio traits are useful heuristics in clinical practice, but introduce statistical challenges when used in regression models [29]. GWAS on ratio traits can uncover associations entirely driven by the denominator, and it is not possible to tell from the summary statistics of a ratio analysis alone which associations are likely denominator-driven. While many associations detected in ratio GWAS do in fact affect the numerator, it is worth scrutinizing the results of ratio GWAS to understand how those associations arose. We recommend comparing the effect sizes from ratio GWAS with those from unadjusted (marginal) GWAS of the numerator and denominator, and from the adjusted model. Variants not strongly associated with the denominator that have similar effect sizes in the unadjusted and adjusted analyses are likely numerator-driven. For such variants, reduction in residual variation by conditioning on the denominator can increase power. Conversely, variants strongly associated with the denominator that display markedly different effect sizes in the adjusted and unadjusted analyses may be denominator-driven.

Pleiotropic effects on the numerators and denominators of ratio traits appear widespread. Pleiotropy can arise either in the vertical sense, as when the effect of a variant on the numerator is mediated by the denominator or vice versa; or in the horizontal sense, as when the variants affects the numerator and denominator through distinct pathways. In the survey of discrepancies between the ratio and adjusted models, 44% of loci identified in the ratio model were detected by the adjusted model and 68% of the loci identified in the adjusted model were detected by the ratio model (**Table S1**). Moreover, the genetic correlation between the numerator and denominator traits was often substantial (**Table S2**). Thus, it is not the case that all associations identified by ratio GWAS are erroneous. Instead, the conclusion is that at least some associations identified by ratio GWAS may be entirely due to the denominator. If the goal of a ratio GWAS is to detect variants associated with the numerator while accounting for the denominator, then an adjusted analysis may be more appropriate. If the goal is to detect pleiotropic variants, i.e. those affecting both the numerator and the denominator, then a multivariate GWAS may outperform either taking the intersection of individual GWAS or performing ratio GWAS [48, 49, 50]. Supplemental Section 1.6 details a multivariate analysis framework as an alternative to ratio analysis. A more recent approach to disentangling pleiotropic from trait-specific effects is genomic structural equation modeling [51].

Among the traits considered in the survey of discrepancies between the ratio and adjusted models, waist-to-hip ratio (WHR) is an outlier. Whereas for other traits, the ratio model detected more associations (as expected), for WHR the adjusted model detected more associations. Moreover, for WHR only, more of the adjusted-only associations than the ratio-only associations were putatively denominator-driven. Possible explanations for this include extensive pleiotropy between waist and hip circumference (using our summary statistics, the genetic correlation was estimated at 0.88 by LDSC; **Table S2**), and the potential for strong environmental effects on both waist and hip circumference (e.g. diet). While unraveling the distinctive behavior of WHR is not the focus of the present work, this does provide an exemplar trait where the adjusted model is likely inappropriate and a multivariate analysis would be preferred.

Recognizing that the purpose of BMI is to provide an index of adiposity that is independent of height, but that the ratio of weight to height^2^ is not always optimal, Stergiakouli *et al* considered performing GWAS on BMI[*x*] = weight*/*height*^x^*, where the exponent *x* was selected in an age-dependent manner to minimize phenotypic correlation between BMI[*x*] and height [52]. Although this is a step in the right direction, this strategy is unlikely to fully resolve the issues with performing GWAS on ratios. Selecting *x* to minimize the phenotypic correlation of BMI[*x*] with height does not guarantee that the final correlation will be exactly zero, and even if zero correlation could be achieved, this does not imply that BMI[*x*] and height will be free of higher-order dependencies. Absent the ability to select *x* ≠ 0 such that BMI[*x*] and height are completely independent, it remains possible for variants to associate with BMI[*x*] via height.

When the denominator is heritable, adjusting for the denominator as a covariate can introduce collider bias [30, 31]. It is worth noting that adjusting for a heritable covariate does not automatically result in collider bias. As shown here and discussed previously [30], there are causal architectures that do not incur collider bias, such as when the effect of background is fully mediated through the heritable covariate. For settings where collider bias is expected, we proposed additionally adjusting for a LOCO-PGS for the heritable covariate. To further control for effects of genetic background on numerator and denominator, this approach could be extended by including a random effect with covariance proportional to the genetic relatedness matrix [53, 45, 54]. When performing GWAS of weight adjusted for height, we found that addition of a LOCO-PGS had little effect on the estimated effect sizes, suggesting that if collider bias is present, it is primarily due to factors other than genetic background. A limitation of the LOCO-PGS approach is that it cannot adjust for collider bias introduced by non-genetic common causes, such as the environment. Methods for addressing collider bias due to potentially unobserved factors include mtCOJO [55], which employs Mendelian randomization, and SlopeHunter [56], which utilizes variants associated with the heritable covariate to estimate a correction factor subsequently applied to all variants. Another limitation is that LOCO-PGS does not, by default, account for collider bias due to variants on the same chromosome. While the bias introduced by variants on the same chromosome will often be minimal, in cases where it is substantial, this bias can be removed by combining LOCO-PGS with conditional GWAS. Specifically, the association model would adjust for the heritable covaraite, its LOCO-PGS, and any variants on the same chromosome with strong effects on the covariate.

With increasingly high-dimensional phenotypic data, such as that provided by metabolomics and proteomics panels, there is a temptation to conduct hypothesis-free association testing on the ratios of all possible pairs of variables [57, 58]. In the metabolic context, a rationale for considering ratios is that the ratio of product to substrate may capture information about binding affinity or reaction rate [11, 58, 19, 59]. While GWAS of ratios may be justified in some situations, we suggest that the scientific hypothesis be carefully elaborated before a ratio phenotype is selected. The fact that the denominator is likely related to the numerator is not sufficient justification for using a ratio phenotype as the relationship between numerator and denominator can be modeled in other ways. If potential collider bias precludes adjusting for the denominator as a covariate, then a multivariate analysis that separately estimates the effect of genotype on the numerator and denominator will typically provide greater interpretability. Notably, the hypotheses that genotype is associated with both the numerator and denominator, or with one or the other, can all be evaluated within a multivariate framework (Supplemental Section 1.6). Performing GWAS on ratios is generally admissible when the denominator is a non-heritable normalizing factor, such as volume in the case of concentration measurements (e.g. HbA1c) or time in the case of certain rates (e.g. 24-hour urinary albumin excretion rate). However, even here it should be considered whether treating the denominator as an offset or covariate is not more appropriate given the other covariates in the model [29].

On the basis of improved power, a recent paper advocated for performing GWAS on log-ratios [21]. Supplemental Section 1.5 contains an extended discussion of performing GWAS on such traits, in which we demonstrate analytically that association with either the numerator or denominator is sufficient for association with the log-ratio. As such, the set of variants targeted by the log-ratio model is the union of the sets of variants affecting the component traits. Direct comparison of power between the log-ratio model and marginal GWAS of the component traits is therefore difficult to justify, because these models target different sets of variants. The log-ratio model, which targets a larger set of variants, will often identify more GWS associations, however the interpretation of which trait these variants affect will be unclear.

Proportions are ratios in which the numerator is contained within the denominator, as in the case of body fat percentage [60]. As ratios, proportions are not exempt from the potential for denominator-driven associations. Taking body fat percentage as an example, suppose total body mass is partitioned into fat mass and lean mass. A genetic variant that increases lean mass but has no direct effect on fat mass will exhibit a negative association with body fat percentage because it increases the denominator (total body mass) via its effect on lean mass.

Although we have scrutinized the practice of using a ratio as the outcome in a regression model, ratios can also enter GWAS as covariates. For example, GWAS of WHR are often adjusted for BMI [41, 61, 62]. When including a ratio as a covariate, it is important to recognize that a ratio is implicitly an interaction; in the case of BMI, between weight and height^−2^ [29]. Including an interaction without the main effects makes it unclear whether an association is genuinely attributable to the interaction versus one of its components; including the main effects in addition to the interaction disentangles these effects [63]. Therefore, in models where adjusting for a ratio is indicated, we follow [29] in suggesting that the components of the ratio also be included as main effects.

In conclusion, GWAS of ratio traits can identify associations that are entirely denominator-driven, and when the rationale for forming the ratio was to adjust for the denominator, such associations may be considered false positives. We recommend reanalyzing ratio GWAS in order to discern whether the associations are attributable to the numerator, the denominator, or both. The practice of considering associations with ratios (e.g. BMI) as conceptually distinct from associations with the components (e.g. height and weight) requires critical reexamination, as variants can associate with the ratio through only one of the components. Downstream analyses based on summary statistics from ratio GWAS (e.g. predictions based on polygenic scores, causal effect estimates based on Mendelian randomization) may also require reconsideration.

## Methods

### Statement on Ethics

This research was conducted using data from the UK Biobank Resource under approved Application Number 51766.

### GWAS catalog analysis

We downloaded all studies in the NHGRI-EBI GWAS Catalog [37] (accessed 07-APR-2023) and searched for the following terms in the “MAPPED TRAIT” field of the studies table: concentration, fraction, index, percent, percentile, proportion, rate, ratio, percentage; BMI, chronic kidney disease, chronic obstructive pulmonary disease, CKD, COPD, and WHR. This returned a collection of 667 traits, which we manually reviewed for evidence of GWAS having been performed on a ratio outcome with a heritable denominator, examining the title, abstract, and study as necessary. The list of 362 ratio traits is provided as Supplementary Data.

### GWAS

GWAS among unrelated White-British subjects from the UK Biobank [32, 33] were performed using PLINK (v1.9) [64]. We filtered to imputed genotypes with minor allele frequency *>* 0.1%, INFO score *>* 0.8, and Hardy-Weinberg equilibrium *P >* 1 *×* 10^−10^. Samples were filtered to those used in the genetic PCA calculation [33]; self-reported “White-British”, “White”, or “Irish” ancestry; and without sex chromosome aneuploidy. For GWAS, samples with phenotype values in the bottom 0.01 or top 99.99 percentiles were removed. To facilitate direct comparison of effect sizes, all models included a standard set of covariates, namely age at recruitment (UKB field: 21022), genetic sex (22001), genotyping array (Axiom vs. BiLEVE; 22000), and the top 20 genetic principal components (22009). Genome-wide significance was declared at the standard threshold of 5 *×* 10^−8^ [34].

### Clumping

Loci independent at *R*^2^ *≤* 0.1 and genome-wide significant (GWS) at *P ≤* 5 *×* 10^−8^ loci were identified by applying PLINK’s clumping functionality to genome-wide summary statistics with a window size of 250kb. Any reference to independent, GWS loci or clumping refers to clumping with these parameters, unless otherwise noted.

### Effect size correlations

To obtain unbiased estimates of effect size correlation, the full UKB cohort (*N* = 356K) was randomly split into two independent subsets of size *N* = 178K, labeled A and B. Subset A was arbitrarily designated the discovery cohort and subset B the validation cohort. GWAS was performed in both cohorts, and correlations were estimated across cohorts, restricting to variants that were independent and GWS in the discovery cohort. As sensitivity analyses, we explored taking B as the discovery cohort and A as the validation cohort, as well as generating additional random splits. Results from these analyses were qualitatively similar to the results presented.

### Variant group definition

The full set of loci detected by the ratio and adjusted models was partitioned into 3 sets (ratio-only, adjusted-only, or both) based on whether or not the lead variant from each ratio locus was within 250kb of the lead variant for an adjusted locus, and vice versa. Variants having a suggestive association with height (*p <* 1 *×* 10^−6^) were excluded from downstream analysis. Summary statistics from GWAS on obesity-related traits, excluding the UKB cohort where possible, were downloaded from GWAS Catalog, dbGaP, or directly from consortia websites. The summary statistics were lifted over to GRCh38 where needed, and independent, GWS loci were identified by clumping *R*^2^ *≤* 0.5.

### Leave-one-chromosome-out polygenic scores

Leave-one-chromosome-out (LOCO) polygenic scoring was performed within the split-sample framework. First, GWAS for height was performed in each of subsets A and B. Using weights from the independent, GWS variants for height from subset B, a whole-genome polygenic score (PGS) *S* was calculated for each subject in subset A and vice versa for B. Next, for each chromosome *k*, a LOCO-PGS *S_k_* was formed by removing from *S* the contributions of all variants on chromosome *k*. This resulted in 22 LOCO-PGSs per subject, one for each autosome (our GWAS were confined to autosomes). Finally, GWAS was performed, separately for each chromosome, using the association model:

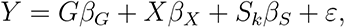

where *G* is genotype, *X* is a set of covariates including height, and *S_k_* is the LOCO-PGS for the chromosome *k* on which *G* resides. For the non-heritable component analysis, height was regressed on the LOCO-PGS to obtain residuals *H_k_*^⊥^, then GWAS was performed, separately for each chromosome, according to the model:

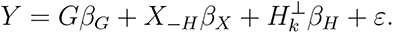

Here the covariate set *X*_−*H*_ now excludes height, and *H_k_*^⊥^ is formed using the *S_k_* for the chromosome on which *G* resides.

## Data Availability

This work used genotypes and phenotypes from the UK Biobank, which are available upon application to the UK Biobank Access Management System (https://www.ukbiobank.ac.uk). Publicly available summary statistics were obtained from the NHGRI-EBI GWAS Catalog (https://www.ebi.ac.uk/gwas/) [37].

## Code Availability

GWAS and clumping were performed using PLINK (v1.9; https://www.cog-genomics.org/ plink/) [64]. Genetic correlations were estimated using LD Score Regression (https://github.com/bulik/ldsc) [36]. Replication code for the simulation studies will be deposited on GitHub.

## Acknowledgements

The authors are grateful to the participants of the UK Biobank, whose data were used with permission.

## Author Contributions

ZRM, HS, SM, and TWS conceived of the project. RD performed theoretical derivations. ZRM, HS, DA, SM, and TWS performed analyses. KS reviewed the list of ratio traits from GWAS catalog. All authors provided scientific input. DK, GDS, DM, and CO provided early feedback and direction on the manuscript. ZRM and TWS wrote the first draft of the manuscript. All authors contributed to critical revision of the final manuscript.

## insitro Research Team Banner and Contribution Statements

All contributors are listed in alphabetical order.

### Downloading, preprocessing, and curation of UK Biobank data

- Francesco Paolo Casale,^1,2,3,4^ Eilon Sharon,^1^ Thomas W. Soare,^1^ James Warren^1^.

### Software engineering

- **Statistical genetics pipeline**: Magdalena Borecka,^1^ Francesco Paolo Casale,^1,2,3,4^ Anna Merkoulovitch,^1^ Colm O’Dushlaine,^1^ Thomas W. Soare,^1^ Paul Sud,^1^ Baris Ungun^1^.
- **redun development**: Robin Betz,^1^ Edward Chee,^1^ Patrick R. Conrad,^1^ Kevin Ford,^1^ Christoph Klein,^1^ Donald Naegely,^1^ Matthew Rasmussen^1^.

