## Supplemental for "Pitfalls in performing genome-wide association studies on ratio traits"

### 1 Supplemental Methods

#### 1.1 Null numerator simulation

Genotypes for $N = 10^4$ independent subjects were simulated at $J = 10^2$ variants in linkage equilibrium. Each variant $G_j$ was standardized to have mean 0 and variance 1. A base denominator phenotype, with heritability $h^2$, was simulated from an infinitesimal model:

$$Y_{h^2} = \sum_{j=1}^{J} G_j \beta_j + \epsilon,$$

where the effect sizes $\beta_j$ were simulated as independent random effects $\beta_j \sim N(0, h^2/J)$, and the residual was simulated as $\epsilon \sim N(0, 1 - h^2)$. In this way:

$$\mathbb{V}(Y_{h^2}) = \sum_{j=1}^{J} \mathbb{V}(G_j)\mathbb{V}(\beta_j) + \mathbb{V}(\epsilon) = \sum_{j=1}^{J} 1 \cdot \left(\frac{h^2}{J}\right) + (1 - h^2) = h^2.$$

Finally the base phenotype $Y$ was shifted by the observed mean $\mu_{\text{obs}}$ and scaled by the observed standard deviation $\sigma_{\text{obs}}$ of height$^2$:

$$H^*_{h^2} = \sigma_{\text{obs}} Y_{h^2} + \mu_{\text{obs}}.$$

Across simulations, the mean and SD of BMI were 27.8 and 6.5 km/m$^2$, which are comparable with the mean and SD of BMI in the full cohort (27.4 and 4.8 km/m$^2$ respectively).

The numerator phenotype was permuted weight $W$, with the mean and variance left at their observed values. For ratio GWAS, $W/H^*_{h^2}$ was regressed on genotype $G_j$, while for the adjusted model, $W$ was regressed on $G_j$ adjusting for $H^*_{h^2}$. The rejection probability at $\alpha = 0.05$ was calculated as:

$$\mathbb{P}(\text{rejection}) = \frac{1}{R} \sum_{r=1}^{R} \sum_{j=1}^{J} \mathbb{I}(p_j \leq 0.05),$$

where $p_j$ is the p-value for association of $G_j$ with the outcome and $R$ is the number of simulation replicates.

#### 1.2 Theoretical analysis of rejection probability versus denominator heritability

Suppose we have the following generative models for the two traits $Y^{(1)}$ (numerator) and $Y^{(2)}$ (denominator), where $Y^{(1)}$ is a null trait, meaning that it has no association with genotype, and $Y^{(2)}$ is affected by $p$ independent causal variants $Z_1, \ldots, Z_p$.

$$
\begin{aligned}
&\text{(Numerator)} \quad Y^{(1)} = \mu_1 1 + \epsilon_1, \quad \epsilon_1 \sim N(0, \sigma_1^2 I) \\
&\text{(Denominator)} \quad Y^{(2)} = \mu_2 1 + \sum_{j=1}^{p} Z_j \alpha_j + \epsilon_2, \\
&\alpha_j \overset{\text{i.i.d}}{\sim} N(0, N\tau^2/p), \quad \epsilon \sim N(0, \sigma^2 I),
\end{aligned}
\tag{1}
$$

where $I$ denotes the identity matrix (of dimension same as the sample-size $N$), $\mu_1, \mu_2$ are fixed scalars, $\epsilon_1, \epsilon_2$ are independent errors, and 1 is the vector of all ones. The genotypes are centered, i.e., $Z_i^\top 1 = 0$, and scaled, i.e., $Z_i^\top Z_i = 1$ for all $i = 1, \ldots, p$. The narrow-sense heritability is then defined as $h^2 = \tau^2/(\sigma^2 + \tau^2) = \tau^2/\sigma_2^2$ where $\sigma_2^2 = \sigma^2 + \tau^2$.

Now, without loss of generality, let us assume we are testing the first causal SNP $G = Z_1$ for association with the ratio trait $Y = Y^{(1)}/Y^{(2)}$ based on the linear model $Y = G\beta + \epsilon_Y$. The squared Wald test statistic for testing $H_0 : \beta = 0$ vs $H_1 : \beta \neq 0$ will be $S^2 = \hat{\beta}^2/\hat{\mathbb{V}}(\hat{\beta})$. Then, the following result provides the asymptotic distribution of $S^2$.

**Result 1.** *Assuming the number of causal SNPs (p) for the denominator trait, and sample-size (N) to be large, asymptotically* $S^2 \sim (1 + \eta)\chi_1^2$, *where,*

$$
\eta = \frac{\left(1 + \frac{3\sigma_2^2}{\mu_2^2}\right)^2 \gamma \sigma_2^2}{\mu_2^2 \left[\frac{\sigma_1^2}{\mu_1^2} + \frac{\sigma_2^2}{\mu_2^2}\right]} h^2, \quad \gamma = \lim_{N,p\to\infty} \frac{N}{p}.
$$

This result implies that the test will have inflated chi-square statistics for variants that are causal to the denominator trait, and the asymptotic limit of the inflation factor is $(1 + \eta)$.

In addition, when the mean of the numerator trait $\mu_1 \to 0$, then $\eta \to 0$ and therefore the test will have nominal rejection probability when $\mu_1 = 0$, as shown in **Figures 1(b) and S4**.

*Proof.* First, we re-parametrize the generative model for $Y^{(2)}$ as following,

$$Y^{(2)} = \mu_2 1 + G\alpha + \epsilon_2, \qquad \alpha = \alpha_1, \qquad \epsilon_2 = \sum_{j=2}^{p} Z_j \alpha_j + \epsilon,$$

where $\mathbb{E}(\epsilon_2) = 0$ and $\mathbb{V}(\epsilon_2) = (\sigma^2 + \frac{p-1}{p}\tau^2)I \approx \sigma_2^2 I$. Then, using the second order Taylor series expansion around $Y_i^{(1)} = \mu_1$ and $Y_i^{(2)} = \mu_2 + G_i\alpha$ (with the index $i$ denotes the element corresponding to the $i$-th subject), we have,

$$\begin{aligned}
\mu_{Y_i} &= \mathbb{E}(Y_i) \approx \frac{\mu_1}{\mu_2 + G_i\alpha}\left[1 + \frac{\sigma_2^2}{(\mu_2 + G_i\alpha)^2}\right] \\
\sigma_{Y_i}^2 &= \mathbb{V}(Y_i) \approx \frac{\mu_1^2}{(\mu_2 + G_i\alpha)^2}\left[\frac{\sigma_1^2}{\mu_1^2} + \frac{\sigma_2^2}{(\mu_2 + G_i\alpha)^2}\right].
\end{aligned}$$

Assuming $G_i\alpha = O_p(1/\sqrt{p})$ where $p$ is large, we can further apply Taylor series expansion of $\mu_{Y_i}$ and $\sigma_{Y_i}^2$ around $G_i\alpha = 0$ and ignore terms that are $O_p(1/p)$,

$$\begin{aligned}
\mu_{Y_i} &\approx \frac{\mu_1}{\mu_2}\left[1 + \frac{\sigma_2^2}{\mu_2^2} - \frac{1}{\mu_2}\left(1 + \frac{3\sigma_2^2}{\mu_2^2}\right) G_i\alpha\right] \\
\sigma_{Y_i}^2 &\approx \frac{\mu_1^2}{\mu_2^2}\left[\frac{\sigma_1^2}{\mu_1^2} + \frac{\sigma_2^2}{\mu_2^2} - \frac{2}{\mu_2}\left(\frac{\sigma_1^2}{\mu_1^2} + \frac{2\sigma_2^2}{\mu_2^2}\right) G_i\alpha\right]
\end{aligned} \tag{2}$$

Since $G$ is centered and scaled ($G^\top G = 1$), a linear regression of $Y$ on $G$ will yield the effect-size estimate and the estimate of its variance

$$\hat{\beta} = G^\top Y, \qquad \hat{\mathbb{V}}(\hat{\beta}) = \frac{1}{N-2} Y^\top (I - P_{[1:G]})Y,$$

where $P_A$ denotes the projection matrix of the matrix $A$, and $[1 : G]$ denotes the matrix that includes as columns the ones vector and the genotype vector $G$.

Let us denote $\Sigma_Y$ to be a diagonal matrix with $i$-th diagonal element equal to $\sigma_{Y_i}^2$. We can then show that,

$$
\begin{aligned}
\hat{\mathbb{V}}(\hat{\beta}) \xrightarrow{p} \lim_{N\to\infty} E\left(\hat{\mathbb{V}}(\hat{\beta})\right) &= \lim_{N\to\infty} \frac{1}{N-2}\left[\mathrm{tr}((I-P_{[1:G]})\Sigma_Y) + \mu_Y^\top (I-P_{[1:G]})\mu_Y\right] \\
&= \lim_{N\to\infty} \frac{1}{N-2}\mathrm{tr}((I-P_{[1:G]})\Sigma_Y) \\
&= \lim_{N\to\infty} \frac{1}{N-2}\mathrm{tr}\left[(I-\frac{1}{N}11^\top - GG^\top)\Sigma_Y\right] \\
&= \lim_{N\to\infty} \frac{1}{N}\sum_{i=1}^N \sigma_{Y_i}^2 = \frac{\mu_1^2}{\mu_2^2}\left[\frac{\sigma_1^2}{\mu_1^2}+\frac{\sigma_2^2}{\mu_2^2}\right],
\end{aligned}
$$

The second step follows from the fact that $\mu_Y$ can be written as $\mu_Y = a + bG$ for some constants $a$ and $b$ (2), and hence $(I - P_{[1:G]})\mu_Y = 0$. Now, using (1) and (2),

$$
\begin{aligned}
\mathbb{E}(\hat{\beta}) &= \mathbb{E}\left(\mathbb{E}(\hat{\beta}|\alpha)\right) = \mathbb{E}\left[-\frac{\mu_1}{\mu_2^2}\left(1+\frac{3\sigma_2^2}{\mu_2^2}\right)\alpha\right] = 0, \\
\mathbb{V}(\hat{\beta}) &= \mathbb{E}\left(\mathbb{V}(\hat{\beta}|\alpha)\right) + \mathbb{V}\left(\mathbb{E}(\hat{\beta}|\alpha)\right) \\
&= \mathbb{E}\left[\sum_{i=1}^N G_i^2\sigma_{Y_i}^2\right] + \mathbb{V}\left[-\frac{\mu_1}{\mu_2^2}\left(1+\frac{3\sigma_2^2}{\mu_2^2}\right)\alpha\right] \\
&= \frac{\mu_1^2}{\mu_2^2}\left[\frac{\sigma_1^2}{\mu_1^2}+\frac{\sigma_2^2}{\mu_2^2}\right] - \mathbb{E}\left[\frac{2\mu_1^2}{\mu_2^3}\left(\frac{\sigma_1^2}{\mu_1^2}+\frac{2\sigma_2^2}{\mu_2^2}\right)\alpha\sum_{i]1}^N G_i^3\right] + \frac{\mu_1^2}{\mu_2^4}\left(1+\frac{3\sigma_2^2}{\mu_2^2}\right)^2\frac{N\tau^2}{p} \\
&= \frac{\mu_1^2}{\mu_2^2}\left[\frac{\sigma_1^2}{\mu_1^2}+\frac{\sigma_2^2}{\mu_2^2}\right]\left(1+\frac{\left(1+\frac{3\sigma_2^2}{\mu_2^2}\right)^2\frac{N\tau^2}{p}}{\mu_2^2\left[\frac{\sigma_1^2}{\mu_1^2}+\frac{\sigma_2^2}{\mu_2^2}\right]}\right).
\end{aligned}
$$

Therefore, $|\hat{\mathbb{V}}(\hat{\beta}) - \mathbb{V}(\hat{\beta})/(1+\eta)| \xrightarrow{p} 0$. By replacing the $\hat{\mathbb{V}}(\hat{\beta})$ with its in-probability limit in the expression of the Z-score, we complete the proof,

$$
S^2 = \frac{\hat{\beta}^2}{\hat{\mathbb{V}}(\hat{\beta})} \approx \frac{\hat{\beta}^2}{\mathbb{V}(\hat{\beta})}(1+\eta) \sim (1+\eta)\chi^2 \quad \text{asymptotically.}
$$

□

#### 1.3 Effect size correlations

Consider performing GWAS for two related phenotypes $Y_1$ and $Y_2$ via the following models:

$$Y_1 = G\alpha_G + X_1\alpha_X + \epsilon_1, \qquad Y_2 = G\beta_G + X_2\beta_X + \epsilon_2. \tag{3}$$

Here $G$ is genotype at a particular variant, $X_1$ and $X_2$ are two sets of covariates, not necessarily including the same variables, and $\epsilon_1$ and $\epsilon_2$ are residuals. To simplify subsequent analysis, we can remove dependence on covariates by regressing $X_k$ out of the association model for $Y_k$, $k \in \{1, 2\}$. Define the projection matrices:

$$Q_1 = I - X_1(X_1^T X_1)^{-1}X_1^T, \qquad Q_2 = I - X_2(X_2^T X_2)^{-1}X_2^T.$$

The genetic effect sizes $(\alpha_G, \beta_G)$ from (3) are consistently estimated via the models:

$$e_{Y_1} = e_{G_1}\alpha_G + \varepsilon_1, \qquad e_{Y_2} = e_{G_2}\beta_G + \varepsilon_2, \tag{4}$$

where $e_{Y_k} = Q_k Y_k$ and $e_{G_k} = Q_k G$. That is, $e_{Y_k}$ is the residual after regressing $Y_k$ on $X_k$ and $e_{G_k}$ is the residual after regressing $G$ on $X_k$. Now, the ordinary least squares (OLS) estimates of $\alpha_G$ and $\beta_G$ are:

$$\hat{\alpha}_G = (e_{G_1}^T e_{G_1})^{-1} e_{G_1}^T e_{Y_1}, \qquad \hat{\beta}_G = (e_{G_2}^T e_{G_2})^{-1} e_{G_2}^T e_{Y_2}. \tag{5}$$

Upon substitution of (4) into (5):

$$\hat{\alpha}_G = \alpha_G + (e_{G_1}^T e_{G_1})^{-1} e_{G_1}^T \varepsilon_1, \qquad \hat{\beta}_G = \beta_G + (e_{G_2}^T e_{G_2})^{-1} e_{G_2}^T \varepsilon_2$$

Taking the covariance of the estimated effect sizes:

$$\mathbb{C}(\hat{\alpha}_G, \hat{\beta}_G) = \mathbb{C}(\alpha_G, \beta_G) + (e_{G_1}^T e_{G_1})^{-1} e_{G_1}^T \mathbb{C}(\varepsilon_1, \varepsilon_2) e_{G_2} (e_{G_2}^T e_{G_2})^{-1}.$$

Observe that covariance between the estimated effect sizes has two sources: covariance of the true effect sizes $\mathbb{C}(\alpha_G, \beta_G)$ plus a term that depends on the covariance of residuals $\mathbb{C}(\varepsilon_1, \varepsilon_2)$.

When $\hat{\alpha}_G$ and $\hat{\beta}_G$ are estimated using independent subjects, then $Y_1$ and $Y_2$ are independent, as are the residuals $\epsilon_1$ and $\epsilon_2$ in (3), and by extension the residuals $\varepsilon_1$ and $\varepsilon_2$ in the reduced model (4). Thus, in the setting of independent data sets $\mathbb{C}(\hat{\alpha}_G, \hat{\beta}_G)$ provides an unbiased estimate of the true effect size covariance $\mathbb{C}(\alpha_G, \beta_G)$. However, when $Y_1$ and $Y_2$ are measured on the same subjects, then in general $\epsilon_1$ and $\epsilon_2$ are dependent, and $\mathbb{C}(\varepsilon_1, \varepsilon_2) \neq 0$. Thus, when the same data are utilized to estimate both $\hat{\alpha}_G$ and $\hat{\beta}_G$, $\mathbb{C}(\hat{\alpha}_G, \hat{\beta}_G)$ is biased as an estimate of $\mathbb{C}(\alpha_G, \beta_G)$ with bias depending on $\mathbb{C}(\varepsilon_1, \varepsilon_2)$, which in turn depends on the covariance between the initial phenotypes, $Y_1$ and $Y_2$.

#### 1.4 Leave-one-chromosome-out polygenic scoring

In this section, we analytically show the following:

1. The adjusted model $Y \sim G + X$ is subject to bias when there exist background variants that affect both $X$ and $Y$.
2. This bias can be removed by adjusting for a polygenic score (PGS) for $X$ constructed from variants not in linkage disequilibrium (LD) with the variant of interest $G$.
3. Including $G$ and variants in LD with $G$ in the PGS $S$ results in bias.

##### 1.4.1 Generative model

Consider the following generative models for the numerator ($Y$) and the denominator ($X$) traits,

$$
\begin{aligned}
Y &= G\beta_{GY} + \sum Z_i \beta_{Yi} + \epsilon_Y, \\
X &= G\beta_{GX} + \sum Z_i \beta_{Xi} + \epsilon_X.
\end{aligned}
\tag{6}
$$

Here, $G$ is the variant of interest, $Z_i$-s are the other variants affecting the numerator and/or denominator, and $\epsilon_X \sim N(0, \sigma_X^2 I_N), \epsilon_Y \sim N(0, \sigma_Y^2 I_N)$ are independent errors. For notational simplicity, we assume $G$ and $Z_i$-s are all centered and scaled such that $G^\top G = Z_i^\top Z_i = N$

for all $i$. We further assume that:

$$Z_i = G\alpha_i + \epsilon_i.$$

The parameters $\beta_{Yi}, \beta_{Xi}$, and $\alpha_i$ can assume the value zero. If $\alpha_i = 0$, the variant $Z_i$ is not in LD with $G$, and $Z_i = \epsilon_i$. If $\beta_{Xi} = 0$, then the variant $Z_i$ is not causal to $X$, and similarly for $\beta_{Yi}$.

##### 1.4.2 Adjusted analysis

Suppose we fit the adjusted model $Y \sim G + X$:

$$Y = 1\mu + G\beta + X\gamma + \varepsilon, \tag{7}$$

which is a mis-specified model compared to the true generative model in (6). Let us denote $\tilde{X} = [1 : X]$ to be the covariate matrix $X$ augmented to include the intercept, and $P_A = A(A^\top A)^{-1}A^\top$ to be the matrix for orthogonal projection onto $A$. Then, the effect-size estimate of $G$ from (7) will be,

$$\begin{aligned}
\hat{\beta} &= \left(G^\top(I - P_{\tilde{X}})G\right)^{-1} G^\top(I - P_{\tilde{X}})Y \\
&= \left(G^\top(I - P_{\tilde{X}})G\right)^{-1} G^\top(I - P_{\tilde{X}}) \left(\mu_Y 1 + G\beta_{GY} + \sum Z_i\beta_{Yi} + \epsilon_Y\right) \\
&= \beta_{GY} + \sum \beta_{Yi} \left(G^\top(I - P_{\tilde{X}})G\right)^{-1} G^\top(I - P_{\tilde{X}})Z_i \\
&\qquad + \left(G^\top(I - P_{\tilde{X}})G\right)^{-1} G^\top(I - P_{\tilde{X}})\epsilon_Y.
\end{aligned}$$

Taking expectations,

$$\begin{aligned}
\mathbb{E}(\hat{\beta}) &= \beta_{GY} + \sum \beta_{Yi}\mathbb{E}\left[\left(G^\top(I - P_{\tilde{X}})G\right)^{-1} G^\top(I - P_{\tilde{X}})Z_i\right] \\
&= \beta_{GY} + \sum \beta_{Yi}\mathbb{E}(\hat{\beta}_{\{G:Z_i\sim G+X\}})
\end{aligned}$$

Here, $\hat{\beta}_{\{G:Z_i\sim X+G\}} = \left(G^\top(I - P_{\tilde{X}})G\right)^{-1} G^\top(I - P_{\tilde{X}})Z_i$ is the estimated coefficient for $G$ in the model $Z_i \sim X + G$. The following derivation closely follows Aschard *et al* [1]. First, note that:

$$(I - P_{\tilde{X}})G = (I - P_{[1:X]})G = (I - P_1 - P_{(I-P_1)X})G = (I - P_{X^{(c)}})G,$$

where $X^{(c)} = (I - P_1)X$ is the centered denominator. Next, we have,

$$G^\top (I - P_{\tilde{X}})G = N - \frac{(G^\top X^{(c)})^2}{X^{(c)\top} X^{(c)}} = N \left(1 - \frac{\hat{\beta}_{GX}^2}{v_X}\right),$$

where $\hat{\beta}_{GX}$ is the coefficient estimate of $G$ in the regression of $X$ on $G$, and $v_X = N^{-1} X^{(c)\top} X^{(c)}$ the sample variance of $X$. Based on our generative models, $\hat{\beta}_{GX}$ is unbiased to $\beta_{GX}$. Next,

$$\begin{aligned} G^\top (I - P_{\tilde{X}})Z_i &= G^\top Z_i - \frac{(G^\top X^{(c)})(X^{(c)\top} Z_i)}{X^{(c)\top} X^{(c)}} \\ &= N \left(\hat{\alpha}_i - \frac{\hat{\beta}_{GX} \hat{\rho}_{X,Z_i}}{\sqrt{v_X}}\right), \end{aligned}$$

where $\rho_{X,Z_i}$ is the sample correlation of $X$ and $Z_i$. Therefore,

$$\hat{\beta}_{\{G:Z_i \sim G+X\}} = \frac{\hat{\alpha}_i - \frac{\hat{\beta}_{GX} \hat{\rho}_{X,Z_i}}{\sqrt{v_X}}}{1 - \frac{\hat{\beta}_{GX}^2}{v_X}} \approx \hat{\alpha}_i - \frac{\hat{\beta}_{GX} \hat{\rho}_{X,Z_i}}{\sqrt{v_X}},$$

assuming $\hat{\beta}_{GX}/v_X \ll 1$ as is typical for human traits (this assumption follows [1]). For large sample-sizes, we can replace the sample estimates to their corresponding limits in the population to obtain,

$$\begin{aligned} \mathbb{E}(\hat{\beta}_{\{G:Z_i \sim G+X\}}) &\approx \alpha_i - \frac{\beta_{GX} \rho_{X,Z_i}}{\sqrt{V_X}}, \\ V_X &= \mathbb{E}\left(\frac{1}{N} X^\top (I - P_1) X\right), \\ \rho_{X,Z_i} &= V_X^{-1/2} \mathbb{E}\left(\frac{1}{N} X^\top (I - P_1) Z_i\right), \end{aligned}$$

where $\rho_{X,Z_i}$ is the marginal correlation between $X$ and $Z_i$ and $V_X$ is the marginal variance of $X$ marginalized over all $Z_i$-s, $G$, and $\epsilon_X$. Therefore,

$$\mathbb{E}(\hat{\beta}) \approx \beta_{GY} + \sum \beta_{Yi}(\alpha_i - V_X^{-1/2}\rho_{X,Z_i}\beta_{GX}). \tag{8}$$

In equation (8), $\beta_{GY}$ is the effect size of interest and the summation is a bias term. If $\beta_{Yi} = 0$, meaning the background variants do not affect the numerator trait, then the bias vanishes. However, when $\beta_{Yi} \neq 0$, bias is expected if $Z_i$ is in LD with $G$, such that $\alpha_i \neq 0$, or if

i. $Z_i$ has an effect on $X$, such that $\rho_{X,Z_i} \neq 0$, and

ii. $G$ has an effect on $X$, such that $\beta_{GX} \neq 0$.

##### 1.4.3 Model with a polygenic score

Lets $S$ denote a PGS for the denominator trait $X$. Suppose we update the association model to $Y \sim G + X + S$:

$$Y = 1\mu + G\beta + X\gamma + S\delta + \varepsilon. \tag{9}$$

Then, the effect-size estimate of $G$ from (9) will be:

$$\begin{aligned}
\hat{\beta} &= \left(G^\top(I - P_{[\tilde{X}:S]})G\right)^{-1} G^\top(I - P_{[\tilde{X}:S]})Y \\
&= \beta_{GY} + \sum \beta_{Yi} \left(G^\top(I - P_{[\tilde{X}:S]})G\right)^{-1} G^\top(I - P_{[\tilde{X}:S]})Z_i \\
&\quad + \left(G^\top(I - P_{[\tilde{X}:S]})G\right)^{-1} G^\top(I - P_{[\tilde{X}:S]})\epsilon_Y.
\end{aligned}$$

Taking expectations,

$$\begin{aligned}
\mathbb{E}(\hat{\beta}) &= \beta_{GY} + \sum \beta_{Yi}\mathbb{E}\left[\left(G^\top(I - P_{[\tilde{X}:S]})G\right)^{-1} G^\top(I - P_{[\tilde{X}:S]})Z_i\right] \\
&= \beta_{GY} + \sum \beta_{Yi}\mathbb{E}(\hat{\beta}_{\{G:Z_i \sim G+X+S\}})
\end{aligned}$$

where $\hat{\beta}_{\{G:Z_i \sim G+X+S\}}$ is the coefficient on $G$ from the model $Z_i \sim G+X+S$. In the following, we write:

$$
\begin{aligned}
(I - P_{[\tilde{X}:S]})G &= (I - P_1 - P_{(I-P_1)S} - P_{(I-P_{[1:S]})X})G \\
&= (I - P_{S^{(c)}} - P_{X^{(r)}})G,
\end{aligned}
$$

where $S^{(c)}$ is the centered PGS, and $X^{(r)}$ is the residual after least squares regression of $X$ on $S$, $X \sim S$.

###### 1.4.4 Case 1: PGS constructed only using variants not in LD with $G$

Suppose $S$ is constructed only using variants that are not in LD with $G$. Here, $S^{(c)}$ and $G$ are independent, thus $P_{S^{(c)}}G \xrightarrow{p} 0$. Therefore,

$$
\begin{aligned}
\hat{\beta}_{\{G:Z_i \sim G+X+S\}} &= \left(G^\top (I - P_{[\tilde{X}:S]})G\right)^{-1} G^\top (I - P_{[\tilde{X}:S]})Z_i \\
&\approx \left(G^\top (I - P_{X^{(r)}})G\right)^{-1} G^\top (I - P_{X^{(r)}})Z_i \\
&= \hat{\beta}_{\{G:Z_i \sim G+X^{(r)}\}}
\end{aligned}
$$

We can analyze the above expression as we did for the adjusted model, with the only difference being $X$ is replaced by $X^{(r)}$:

$$
\mathbb{E}(\hat{\beta}_{\{G:Z_i \sim G+X^{(r)}\}}) \approx \alpha_i - \frac{\beta_{GX^{(r)}} \rho_{X^{(r)},Z_i}}{\sqrt{V_X^{(r)}}}.
$$

Now if the $i$th variant is not in LD with $G$, then $\alpha_i = 0$ and $\rho_{X^{(r)},Z_i} \xrightarrow{p} 0$ because $X^{(r)}$ is the residual of $X$ having regressed out the effects of variants not in LD with $G$ (including $Z_i$). Therefore, using a PGS $S$ for $X$ constructed only from variants not in LD with $G$ can remove the heritable covariate bias arising from variants not in LD with $G$.

##### 1.4.5 Case 2: global PGS constructed based on all variants

Suppose instead $S$ is constructed from all variants, including $G$ and variants in LD with $G$. In this case $S^{(c)}$ is no longer uncorrelated with $G$. Then,

$$\begin{aligned}\hat{\beta}_{\{G:Z_i\sim G+X+S\}} &= \left(G^\top(I - P_{[\tilde{X}:S]})G\right)^{-1} G^\top(I - P_{[\tilde{X}:S]})Z_i \\ &\approx \left(G^\top(I - P_{S^{(c)}} - P_{X^{(r)}})G\right)^{-1} G^\top(I - P_{S^{(c)}} - P_{X^{(r)}})Z_i.\end{aligned}$$

Following a similar derivation as before, we can obtain the individual bias contribution terms,

$$\mathbb{E}(\hat{\beta}_{\{G:Z_i\sim G+X+S\}}) \approx \alpha_i - V_X^{(r)-1/2}\beta_{GX^{(r)}}\rho_{X^{(r)},Z_i} - V_S^{(c)-1/2}\beta_{GS^{(c)}}\rho_{S^{(c)},Z_i}.$$

In this case, even when the $i$-th variant is not in LD with $G$, such that $\alpha_i = 0$ and $\rho_{X^{(r)},Z_i} \xrightarrow{p} 0$, there remains a bias term:

$$\mathbb{E}(\hat{\beta}_{\{G:Z_i\sim G+X+S\}}) \approx -V_S^{(c)-1/2}\beta_{GS^{(c)}}\rho_{S^{(c)},Z_i}.$$

$\beta_{GS^{(c)}}$ and $\rho_{S^{(c)},Z_i}$ both will be non-ignorable when $G$ and $Z_i$ are included in the PGS $S$.

#### 1.5 Considerations for logarithms of ratios

##### 1.5.1 Unilateral associations

Here we demonstrate in the context of log-ratios that having association with exactly one of the component traits is sufficient for association with the composite trait. Ratios are often composed of two component traits having the same sign, as in the examples of BMI and WHR. Let $Y$ denote the numerator trait, and $X$ the denominator trait. If $Y$ and $X$ always have the same sign, then the range of $Y/X$ is $(0, \infty)$. To improve residual normality, practitioners will often choose to study the logarithm of the ratio $\ln(Y/X)$, whose range is $(-\infty, \infty)$. If we further suppose that $Y$ and $X$ are always positive, then by the properties of logarithms, we can decompose the logarithm of the ratio as the difference of the logarithms: $\ln(Y/X) = \ln(Y) - \ln(X)$. This decomposition makes clear that the composite phenotype $\ln(Y/X)$ is in fact a linear combination of two phenotypes, $\ln(Y)$ and $\ln(X)$. Let $G$ denote genotype, and $Z$ a set of covariates. Consider the following association models for the log-scale numerator and denominator traits:

$$\begin{aligned} \mathbb{E}\{\ln(Y)|G, Z\} &= G\beta_G + Z\beta_Z, \\ \mathbb{E}\{\ln(X)|G, Z\} &= G\alpha_G + Z\alpha_Z. \end{aligned} \tag{10}$$

Note that we have written the regression models as conditional expectations, making clear the variables being conditioned on. Taking the difference of the equations in (10) gives:

$$\mathbb{E}\{\ln(Y)|G, Z\} - \mathbb{E}\{\ln(X)|G, Z\} = G(\beta_G - \alpha_G) + Z(\beta_Z - \alpha_Z). \tag{11}$$

The difference on the left-hand side is in fact a regression model for the log-ratio:

$$\mathbb{E}\{\ln(Y)|G, Z\} - \mathbb{E}\{\ln(X)|G, Z\} = \mathbb{E}\{\ln(Y) - \ln(X)|G, Z\} = \mathbb{E}\{\ln(Y/X)|G, Z\},$$

where the first equality is due to the linearity of expectation, and the second to the properties of logarithms. Thus, (11) is equivalent to:

$$\mathbb{E}\{\ln(Y/X)|G, Z\} = G(\beta_G - \alpha_G) + Z(\beta_Z - \alpha_Z) \tag{12}$$

Comparison of (12) to (10) makes clear that an association between $G$ and *either* $\ln(Y)$ or $\ln(X)$ is sufficient for the existence of an association between $G$ and the log-ratio. In detail, the coefficient on $G$ in the log-ratio model, namely $(\beta_G - \alpha_G)$, is non-zero in *any* of the following events:

i. $G$ is associated with the numerator only: $\beta_G \neq 0$ and $\alpha_G = 0$.

ii. $G$ is associated with the denominator only: $\beta_G = 0$ and $\alpha_G \neq 0$.

iii. $G$ is associated with both numerator and denominator, and the effect sizes are not identical: $\beta_G \neq 0$, $\alpha_G \neq 0$, $\beta_G \neq \alpha_G$.

An important implication is that the set of variants that can, with sufficient power, be detected by GWAS of the log ratio is the union of the set of variants associated with the numerator plus the set of variants associated with the denominator (minus the set of variants that have exactly the same effect on both). While we have considered the case of log-ratios for clarity, this principle holds more generally: GWAS of a composite trait formed as some function $g(X, Y)$ of two component traits is susceptible to identifying variants associated with either of the component traits.

##### 1.5.2 Conditioning

Suppose again that $Y$ and $X$ are strictly positive numerator and denominator traits, and consider the association model:

$$\ln\left(\frac{Y}{X}\right) = G\beta_G + Z\beta_Z + \epsilon,$$

where $G$ is genotype, $Z$ is a set of covariates, and $\epsilon$ is a residual. By the properties of logarithms, the left-hand side is expressible as:

$$\ln(Y) - \ln(X) = G\beta_G + Z\beta_Z + \epsilon. \tag{13}$$

Now, it may appear that by a simple algebraic operation, namely adding $\ln(X)$ to both sides, (13) can be transformed into the following model:

$$\ln(Y) = \ln(X) + G\beta_G + Z\beta_Z + \epsilon.$$

One may therefore claim that the following adjusted model:

$$\ln(Y) = \ln(X)\beta_X + G\beta_G + Z\beta_Z + \epsilon. \tag{14}$$

is equivalent to (13), with $\beta_X = 1$. This argument is incorrect due to a subtle but important distinction between (13) and (14). By convention, all variables on the left-hand side of a regression model are random, while all variables on the right-hand side, with the exception of the residual, are fixed. This distinction becomes apparent upon taking expectations, clearly identifying the conditioning variables. The regression model in (13) is:

$$\mathbb{E}\{\ln(Y) - \ln(X)|G, Z\} = G\beta_G + Z\beta_Z \tag{15}$$

while that in (14) is:

$$\mathbb{E}\{\ln(Y)|\ln(X), G, Z\} = \ln(X)\beta_X + G\beta_G + Z\beta_Z.. \tag{16}$$

There is a significant difference in interpretation between the effect sizes in (15) and (16). $\beta_G$ in (15) is the expected change in the log-ratio per additional minor allele, holding $Z$ constant. $\beta_G$ in (16) is the expected change in $\ln(Y)$ per additional minor allele, holding $Z$ *and* $\ln(X)$ constant.

##### 1.5.3 Power

It has been argued [2] that ratios improve power as compared to marginal analyses of the numerator and denominator traits. Such a comparison is unfair because, as demonstrated in the subsection on unilateral associations, GWAS of the (log) ratio can detect variants associated with either of the component traits, whereas the marginal GWAS targets variants associated with one specific trait. Moreover, even if we put aside the issue of commensurability and assume that identifying variants associated with either component trait is of interest, there is nothing that uniquely qualifies the ratio for this task. Suppose $Y$ and $X$ are strictly positive numerator and denominator traits, and consider the association models:

$$\ln(Y) = G\beta_G + Z\beta_Z + \epsilon_Y,$$
$$\ln(X) = G\alpha_G + Z\alpha_Z + \epsilon_X,$$

where $G$ is genotype, $Z$ is a set of covariates, and the $\epsilon$s are residuals. Taking the difference gives a model for the log-ratio:

$$\ln(Y) - \ln(X) = \ln\left(\frac{Y}{X}\right) = G(\beta_G - \alpha_G) + Z(\beta_Z - \alpha_Z) + (\epsilon_Y - \epsilon_X).$$

Note that the variance of the residual term for the log-ratio model is:

$$\mathbb{V}(\epsilon_Y - \epsilon_X) = \mathbb{V}(\epsilon_Y) + \mathbb{V}(\epsilon_X) - 2\mathbb{C}(\epsilon_Y, \epsilon_X),$$

where $\mathbb{V}(\cdot)$ denotes the variance and $\mathbb{C}(\cdot, \cdot)$ the covariance. As discussed in [2], power for detecting an association with the log-ratio will generally increase when (i) $\beta_G$ and $\alpha_G$ have opposite signs or (ii) $\epsilon_Y$ and $\epsilon_X$ are positively correlated. Yet it is easy to see that if (i) $\beta_G$ and $\alpha_G$ have the same sign or (ii) $\epsilon_Y$ and $\epsilon_X$ are negatively correlated, then taking the sum of the component models (i.e. modeling the log-product) would improve power:

$$\ln(Y) + \ln(X) = \ln(YX) = G(\beta_G + \alpha_G) + Z(\beta_Z + \alpha_Z) + (\epsilon_Y + \epsilon_X).$$

Moreover, the log-ratio and log-product models are just two special cases of a composite phenotype formed by taking a linear combination of $\ln(Y)$ and $\ln(X)$:

$$g(Y, X) = w_Y \ln(Y) + w_X \ln(X).$$

If the goal is to maximize power, there is no reason to believe that restricting the trait-specific weights ($w_Y$ and $w_X$) to $\{+1, -1\}$ is optimal. In general, selecting these weights adaptively to maximize heritability of the composite phenotype $g(Y, X)$ would be a superior strategy, and considering nonlinear combinations of $Y$ and $X$ may improve power still. While crafting composite phenotypes $g(Y, X)$ to empower genetic discovery is an interesting direction for future research, any composite phenotype formed from multiple heritable components will have the drawback that, without further investigation, we cannot discern which of the component traits drove the association.

#### 1.6 Multivariate modeling

Here we describe a multivariate framework as an alternative to ratio analysis, and show that the hypothesis testing by the log-ratio model in section (1.5.1) is a special case. Let $Y$ denote the numerator trait, and $X$ the denominator trait. Consider the following linear mixed effects model:

$$\begin{pmatrix} Y \\ X \end{pmatrix} \big| (G, Z, \gamma) = \begin{pmatrix} G\beta_G + Z\beta_Z + \gamma \\ G\alpha_G + Z\alpha_Z + \gamma \end{pmatrix} + \begin{pmatrix} \epsilon_Y \\ \epsilon_X \end{pmatrix}.$$

Here $G$ is genotype, $Z$ is a set of covariates, and $\gamma \sim N(0, \sigma_\gamma^2)$ is a per-subject random effect, accounting for within-subject correlation, that is independent of the residuals:

$$\begin{pmatrix} \epsilon_Y \\ \epsilon_X \end{pmatrix} \sim N\left( \begin{pmatrix} 0 \\ 0 \end{pmatrix}, \begin{pmatrix} \sigma_Y^2 & 0 \\ 0 & \sigma_X^2 \end{pmatrix} \right).$$

Marginalizing over $\gamma$ gives the implied model:

$$\begin{pmatrix} Y \\ X \end{pmatrix} \big| (G, Z) \sim N\left( \begin{pmatrix} G\beta_G + Z\beta_Z \\ G\alpha_G + Z\alpha_Z \end{pmatrix}, \begin{pmatrix} \sigma_Y^2 + \sigma_\gamma^2 & \sigma_\gamma^2 \\ \sigma_\gamma^2 & \sigma_X^2 + \sigma_\gamma^2 \end{pmatrix} \right)$$

The model can be generalized by rewriting the covariance matrix as:

$$\boldsymbol{\Sigma} = \begin{pmatrix} \Sigma_{YY} & \Sigma_{YX} \\ \Sigma_{XY} & \Sigma_{XX} \end{pmatrix}$$

and allowing the cross term $\Sigma_{YX} = \Sigma_{XY}$ to assume negative in addition to positive values. We can write the marginal model more compactly by defining some additional notation. Let $\boldsymbol{y} = \binom{Y}{X}$, $W = \begin{bmatrix} G & Z \end{bmatrix}$, $\boldsymbol{W} = \text{diag}(W, W)$, $\beta = \binom{\beta_G}{\beta_Z}$, $\alpha = \binom{\alpha_G}{\alpha_Z}$, $\boldsymbol{\delta} = \binom{\beta}{\alpha}$, then:

$$\boldsymbol{y} | \boldsymbol{W} \sim N\big(\boldsymbol{W}\boldsymbol{\delta}, \boldsymbol{\Sigma}\big).$$

The maximum likelihood estimate (MLE) $\hat{\boldsymbol{\delta}}$ of the regression parameters is asymptotically normal with covariance given by the inverse Fisher information:

$$\begin{pmatrix} \hat{\beta} \\ \hat{\alpha} \end{pmatrix} \dot{\sim} N\left( \begin{pmatrix} \beta \\ \alpha \end{pmatrix}, \begin{pmatrix} \mathcal{I}_{\beta\beta'} & \mathcal{I}_{\beta\alpha'} \\ \mathcal{I}_{\alpha\beta'} & \mathcal{I}_{\alpha\alpha'}, \end{pmatrix}^{-1} \right).$$

The components of the Fisher information are calculated as:

$$\mathcal{I}_{\delta\delta'} = \begin{pmatrix} \mathcal{I}_{\beta\beta'} & \mathcal{I}_{\beta\alpha'} \\ \mathcal{I}_{\alpha\beta'} & \mathcal{I}_{\alpha\alpha'}, \end{pmatrix} = \sum_{i=1}^{n} \begin{pmatrix} W_i^T \Sigma_{YY}^{-} W_i & W_i^T \Sigma_{YX}^{-} W_i \\ W_i^T \Sigma_{XY}^{-} W_i & W_i^T \Sigma_{XX}^{-} W_i \end{pmatrix} = \sum_{i=1}^{n} \begin{pmatrix} \Sigma_{YY}^{-} W_i^T W_i & \Sigma_{YX}^{-} W_i^T W_i \\ \Sigma_{XY}^{-} W_i^T W_i & \Sigma_{XX}^{-} W_i^T W_i \end{pmatrix},$$

where $W_i = \begin{bmatrix} G_i & Z_i \end{bmatrix}$ is the concatenated genotype plus covariate row-vector for subject $i$, and,

$$\Sigma^{-1} = \begin{pmatrix} \Sigma_{YY}^{-} & \Sigma_{YX}^{-} \\ \Sigma_{XY}^{-} & \Sigma_{XX}^{-} \end{pmatrix}.$$

(The final equality uses the fact that the components of $\boldsymbol{\Sigma}^{-1}$ are scalars in a bivariate outcome model.) The marginal hypothesis $H_{0,Y} : \beta_G = 0$ can be evaluated by defining a selection vector $\boldsymbol{e}_Y$ which is equal to 1 where $\boldsymbol{\delta}$ contains $\beta_G$, and 0 elsewhere. Analogously, $H_{0,X} : \alpha_G = 0$ is evaluated by defining selection vector $\boldsymbol{e}_X$ which is 1 where $\boldsymbol{\delta}$ contains $\alpha_G$, and 0 otherwise. With these, $\boldsymbol{e}_Y^T \boldsymbol{\delta} = \beta_G$ and $\boldsymbol{e}_X^T \boldsymbol{\delta} = \alpha_G$. The respective Wald statistics for evaluating $H_{0,Y}$ and $H_{0,X}$ are:

$$T_Y = (\boldsymbol{e}_Y^T \hat{\boldsymbol{\delta}})^T (\boldsymbol{e}_Y^T \mathcal{I}_{\delta\delta'}^{-1} \boldsymbol{e}_Y)^{-1} (\boldsymbol{e}_Y^T \hat{\boldsymbol{\delta}}), \qquad T_X = (\boldsymbol{e}_X^T \hat{\boldsymbol{\delta}})^T (\boldsymbol{e}_X^T \mathcal{I}_{\delta\delta'}^{-1} \boldsymbol{e}_X)^{-1} (\boldsymbol{e}_X^T \hat{\boldsymbol{\delta}})$$

Each Wald statistic is asymptotically $\chi_1^2(0)$ under the null hypothesis. To test the joint hypothesis $H_{0,YX} : (\beta_G = 0) \cap (\alpha_G = 0)$, let $\boldsymbol{E}_{YX} = (\boldsymbol{e}_Y, \boldsymbol{e}_X)$, then the Wald statistic is:

$$T_{YX} = (\boldsymbol{E}_{XY}^T \hat{\boldsymbol{\delta}})^T (\boldsymbol{E}_{XY}^T \mathcal{I}_{\delta\delta'}^{-1} \boldsymbol{E}_{XY})^{-1} (\boldsymbol{E}_{XY}^T \hat{\boldsymbol{\delta}}).$$

Under the null hypothesis, $T_{YX}$ is asymptotically $\chi_2^2(0)$. The framework elaborated here allows for evaluation of the marginal and joint hypotheses using a single model. Moreover, if $Y$ and $X$ are strictly positive and we replace $Y \mapsto \ln(Y)$ and $X \mapsto \ln(X)$, then we can easily test the same hypothesis evaluated by the log-ratio model, namely $H_{0,\text{log-ratio}} : \beta_G - \alpha_G = 0$. In particular, consider the model:

$$\boldsymbol{y}_{\text{log}} | \boldsymbol{W} \sim N(\boldsymbol{W}\boldsymbol{\delta}, \boldsymbol{\Sigma})$$

where $\boldsymbol{y}_{\text{log}} = \binom{\ln(Y)}{\ln(X)}$. $\hat{\boldsymbol{\delta}}$ is estimated by maximum likelihood as before. Define the contrast vector $\boldsymbol{c}$ which is 1 where $\boldsymbol{\delta}$ contains $\beta_G$, $-1$ where $\boldsymbol{\delta}$ contains $\alpha_G$, and otherwise 0, then:

$$T_{\text{log-ratio}} = (\boldsymbol{c}^T\hat{\boldsymbol{\delta}})^T(\boldsymbol{c}^T\mathcal{I}_{\delta\delta'}^{-1}\boldsymbol{c})^{-1}(\boldsymbol{c}^T\hat{\boldsymbol{\delta}})$$

provides a Wald statistic for evaluating $H_{0,\text{log-ratio}}$. We can generalize this statistic to a test of the weighted hypothesis $H_{0,\text{weighted}} : w_Y\beta_G + w_X\alpha_G = 0$ by defining a weight vector $\boldsymbol{w}$ which takes the value $w_Y$ where $\boldsymbol{\delta}$ contains $\beta_G$, $w_X$ where $\boldsymbol{\delta}$ contains $\alpha_G$, and 0 elsewhere. The Wald statistic for this hypothesis is:

$$T_{\text{weighted}} = (\boldsymbol{w}^T\hat{\boldsymbol{\delta}})^T(\boldsymbol{w}^T\mathcal{I}_{\delta\delta'}^{-1}\boldsymbol{w})^{-1}(\boldsymbol{w}^T\hat{\boldsymbol{\delta}}),$$

which is asymptotically $\chi_1^2(0)$ under the null. The $H_{0,\text{log-ratio}}$ is a case of $H_{0,\text{weighted}}$ with $w_Y = 1$ and $w_X = -1$; the log-product hypothesis $H_{0,\text{log-product}} : \beta_G + \alpha_G = 0$ is another.

### 2 Supplemental Results

#### 2.1 Phenotypic landscape

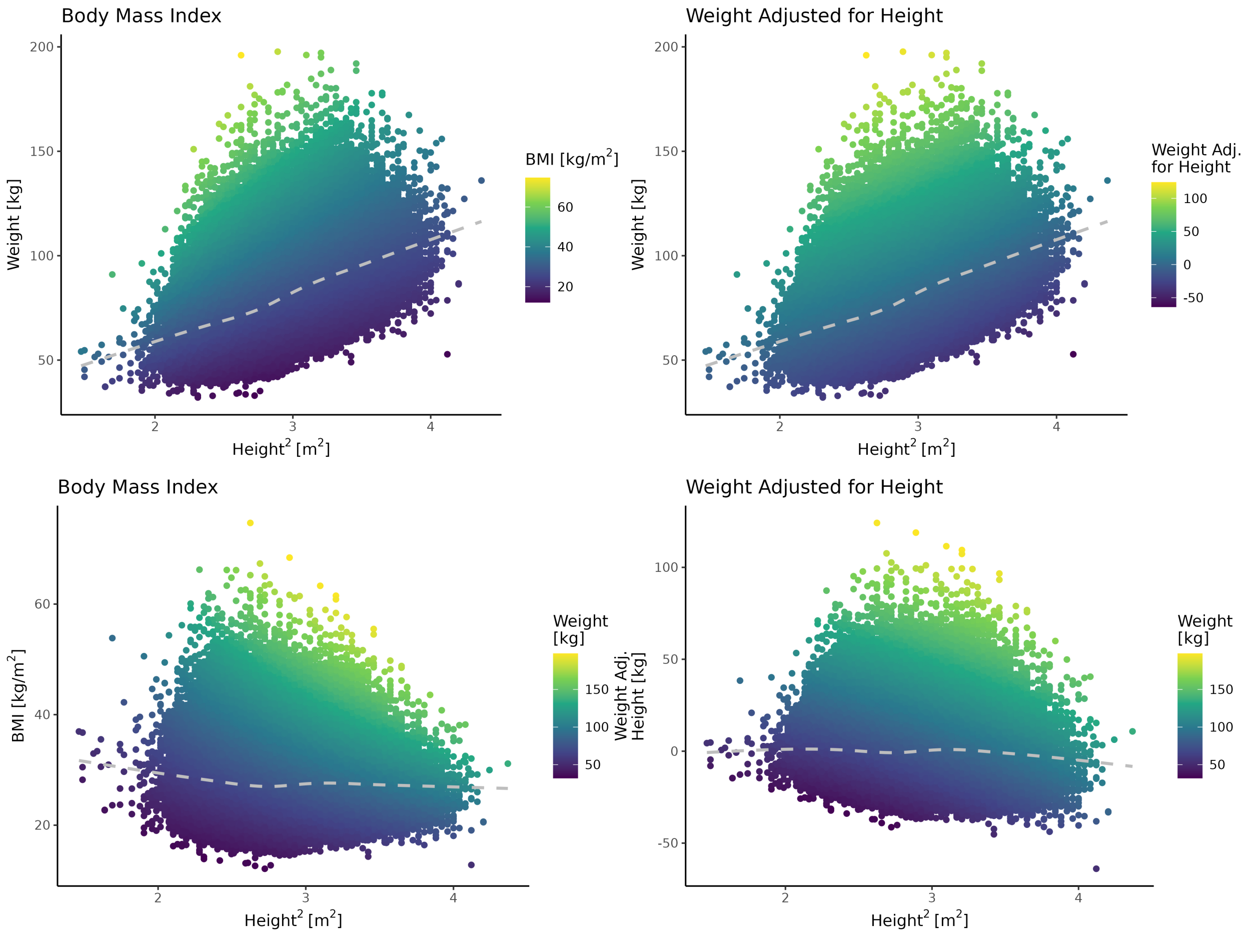

Figure S1: **Association of body mass index and weight adjusted for height with the component traits.** Body mass index (BMI) is defined as the ratio of weight in kilograms (kg) to height in meters$^2$. Weight adjusted for height was obtained by regressing weight on height and height$^2$. The interpretation of adjusted weight is the difference, in kg, of a subject's weight from that expected given their height (and height$^2$), which can be negative. The dashed gray line is a generalized additive model for the trend. Each point is a subject. The association is shown among the full set of unrelated White-British subjects used for GWAS ($N = 356\text{K}$).

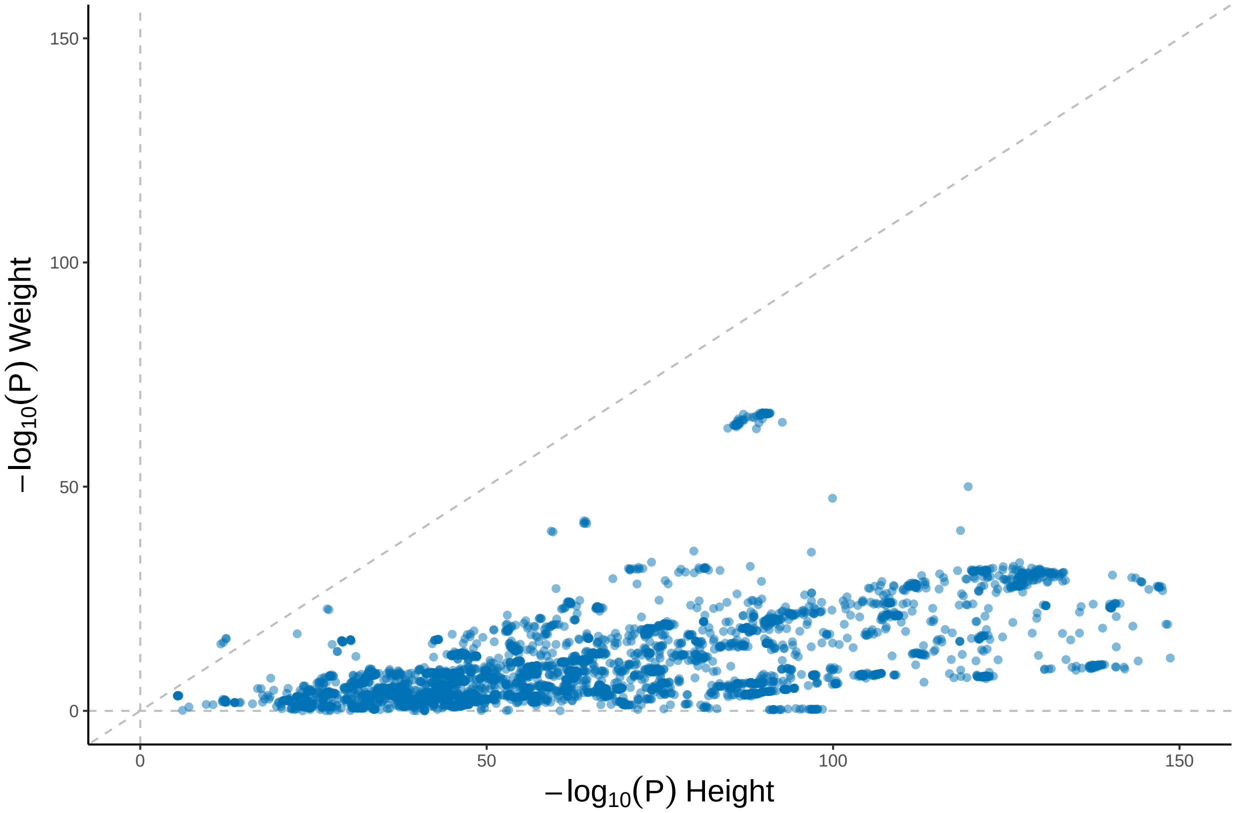

Figure S2: **Associations with weight and height among variants genome-wide significant associations for permuted weight divided by height$^2$.** Shown are the marginal associations with weight and with height among variants genome-wide significant for Permuted Weight/Height$^2$ at $P \leq 5 \times 10^{-8}$. P-values were obtained from GWAS among $N = 356$K unrelated subjects from the UK Biobank. The diagonal line is the identity. Variants below the diagonal exhibit stronger association with height than with weight. Note that some degree of association with height is expected due to pleiotropy and the fact that weight generally increases with height.

#### 2.2 Ratio transformations

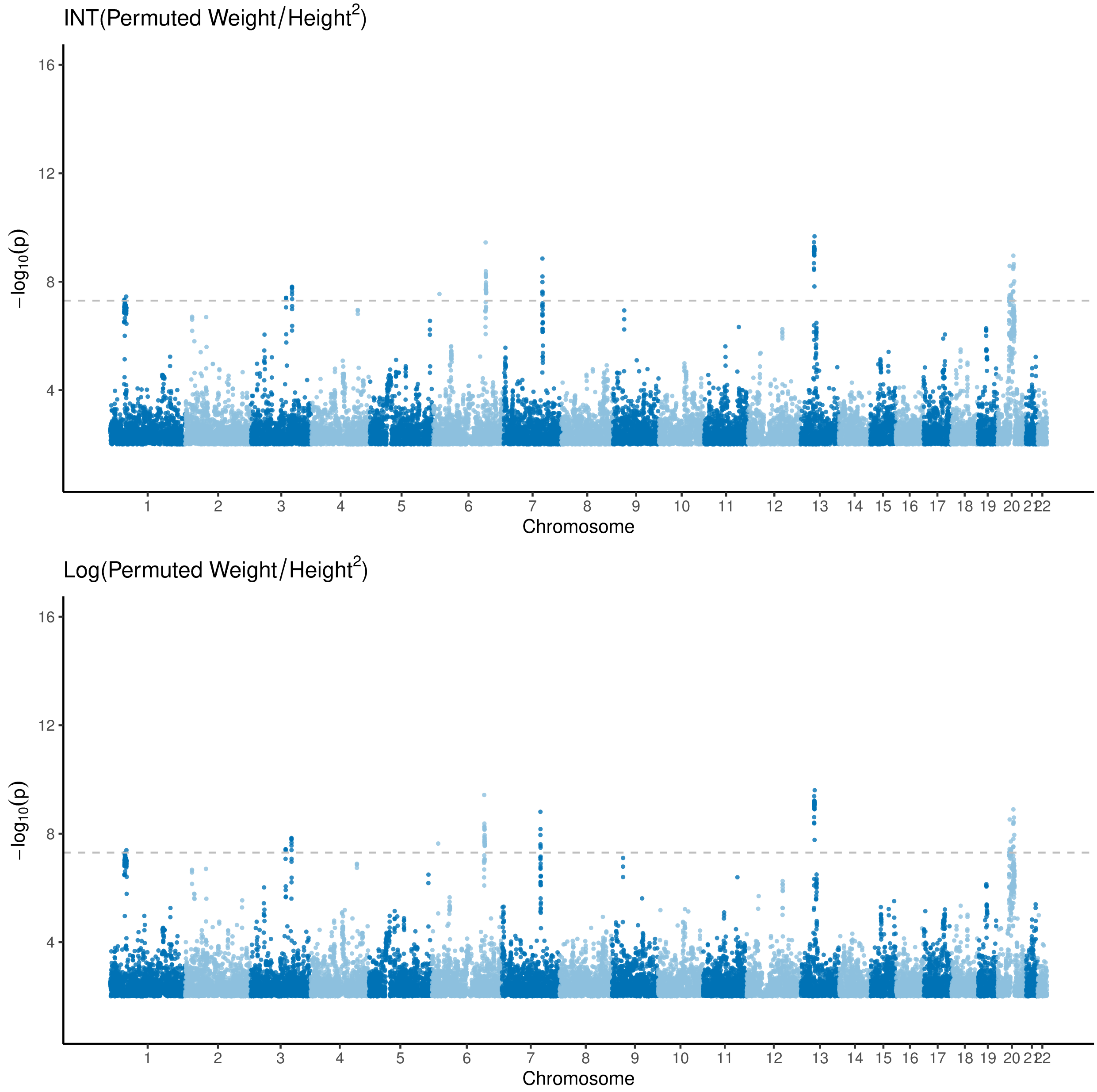

Figure S3: **Ratio GWAS on transformation of permuted weight over height$^2$.** The GWAS were performed among $N = 356\text{K}$ unrelated subjects from the UK Biobank. Each point is a genetic variant. Neither log nor inverse-normal transformation (INT) of the ratio prevent detection of denominator-driven associations.

#### 2.3 Null numerator experiment

Weight was permuted to form a null phenotype $W$, then centered and scaled to have mean 0 and variance 1. The mean of $W$ was shifted by adding a constant $\mu \in \{0, \ldots, 250\}$:

$$W_\mu = W + \mu.$$

$W$ was analyzed either as a ratio, dividing by the observed value of height$^2$ to emulate BMI, or via an adjusted model, conditioning on height and height$^2$. In **Figure S4**, the number of independent genome-wide significant loci is reported as a function of null numerator's mean. Whereas no loci are discovered by the adjusted model, the number of loci detected by the ratio model depends artifactually on the null numerator's mean. This result is not particular to weight; replacing $W$ with independent random draws from the standard normal distribution led to nearly identical results. **Figure S5** shows that the Manhattan plots for observed height and for permuted height, shifted to mean 250, divided by height$^2$ are nearly identical, underscoring that the associations being detected by the ratio model are exactly those associated with the denominator trait.

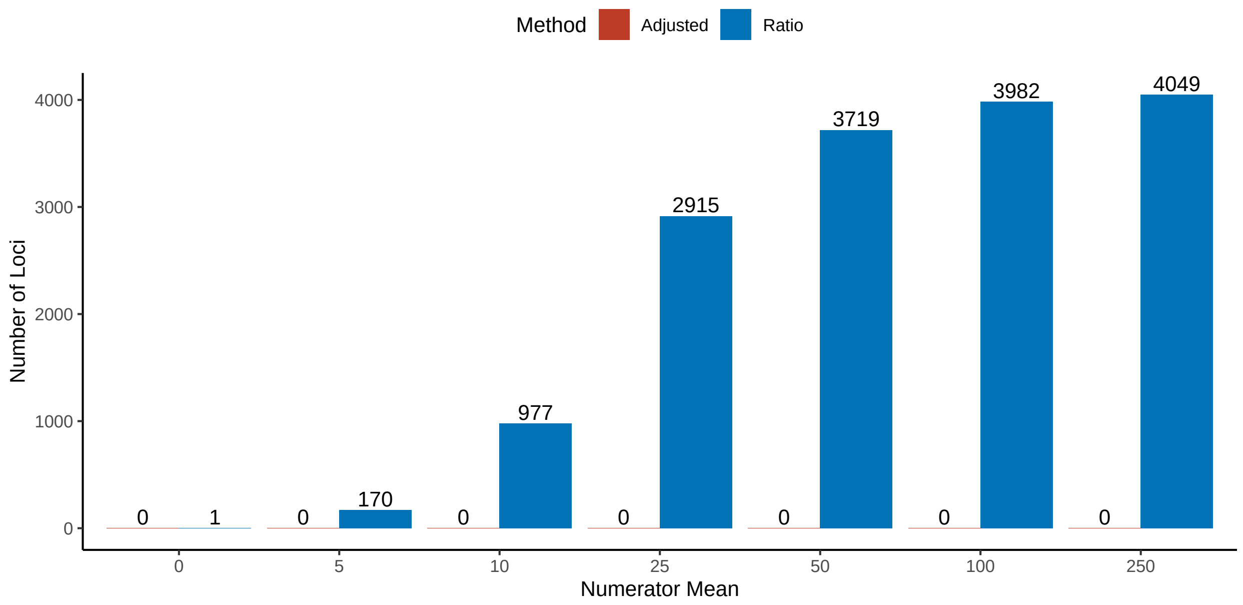

Figure S4: **Artifactual dependence of the number of loci discovered by the ratio model on the mean of the numerator.** A null phenotype was formed by permuting weight, scaling to unit variance, then shifting to mean $\mu$. The null phenotype was either analyzed as a ratio, dividing by the observed value of height$^2$ (to emulate BMI), or via an adjusted model, conditioning on height and height$^2$. Shown are the number of independent ($R^2 \leq 0.1$) genome-wide significant ($P \leq 5 \times 10^{-8}$) loci. The GWAS were performed among $N = 356$K unrelated subjects from the UK Biobank.

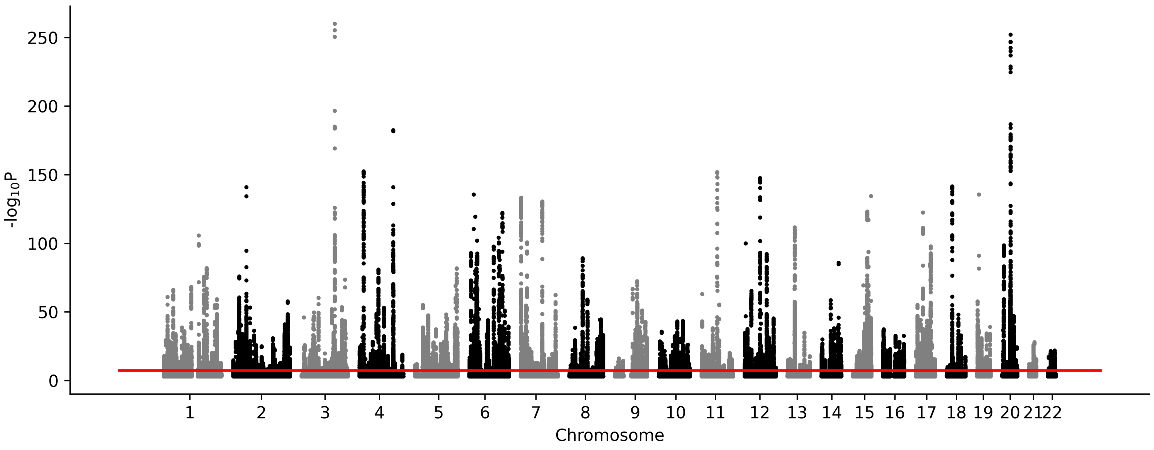

(a) **Height**

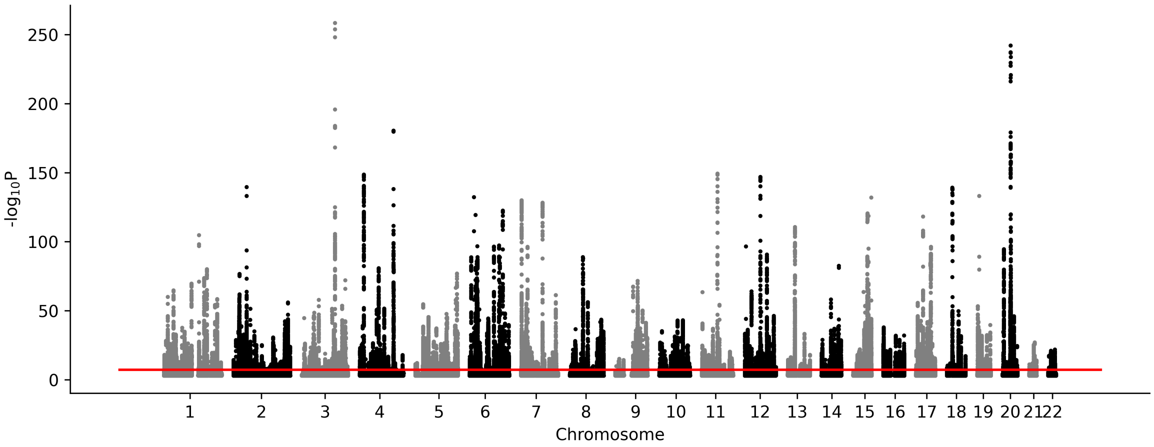

(b) **(Permuted Weight + 250) / Height$^2$**

Figure S5: **Manhattan plots for height and for permuted weighted, shifted to have mean 250, divided by height$^2$.** The GWAS were performed among $N = 356\text{K}$ unrelated subjects from the UK Biobank. Each point is a genetic variant. Because weight is permuted, any associations in the lower plot are attributable to height.

#### 2.4 Environmental correlation simulation

To investigate the impact of environmental correlation on the operating characteristics of the ratio and adjusted models, numerator and denominator traits were simulated for $n = 10^4$ subjects from the following model:

$$Y = \mu_Y + \sigma_Y \varepsilon_Y,$$
$$X = \mu_X + \sigma_X \big(G\alpha_G + \varepsilon_X\big),$$

where $(\mu_Y, \sigma_Y)$ are constants set to match the mean and SD of $Y$ to the empirical mean and SD of weight; $(\mu_X, \sigma_X)$ are constants set to match the mean and SD of $X$ to those of height$^2$; $G$ represents genotype at $J = 10^2$ variants in linkage equilibrium; and $\alpha_G$ was simulated from an infinitesimal model such that the heritable of $X$ was $h^2$. Residuals were simulated from a bivariate normal distribution with correlation $\rho$:

$$\begin{pmatrix} \varepsilon_Y \\ \varepsilon_X \end{pmatrix} \sim N\left\{ \begin{pmatrix} 0 \\ 0 \end{pmatrix}, \begin{pmatrix} 1 & \rho \\ \rho & 1 - h^2 \end{pmatrix} \right\}.$$

Note that under this data generating process, $G$ has no direct effect on $Y$, but $X$ is a collider. That is, there exists background environmental factors that affect both $X$ and $Y$.

**Figures S6 and S7** present the rejection probability and mean $\chi^2$ statistics of the ratio and adjusted models. In the absence of environmental correlation $\rho = 0$, the adjusted model maintains rejection probability near 5% and a mean $\chi^2$ near 1.0, as expected in the absence of association. Meanwhile, the ratio model has rejection probability increasing in the denominator heritability. The rejection probability of the adjusted model increases with the magnitude of the environmental correlation because for $|\rho| > 0$, $X$ is a collider. LOCO-PGS can do nothing to eliminate this collider bias, which is purely environmental in origin. For $|\rho| < 0.5$, the probability of rejection with the adjusted model, due to collider bias, was less than the probability of rejection with the ratio model. At $|\rho| > 0.5$, the rejection probabilities of the two models were similar. Note that the behavior of the adjusted model is symmetric in $\rho$, but that of the ratio model is not. Consistent with the discussion in Supplementary Section 1.5.3, the (log) ratio model has greater rejection in the presence of positively correlated residuals.

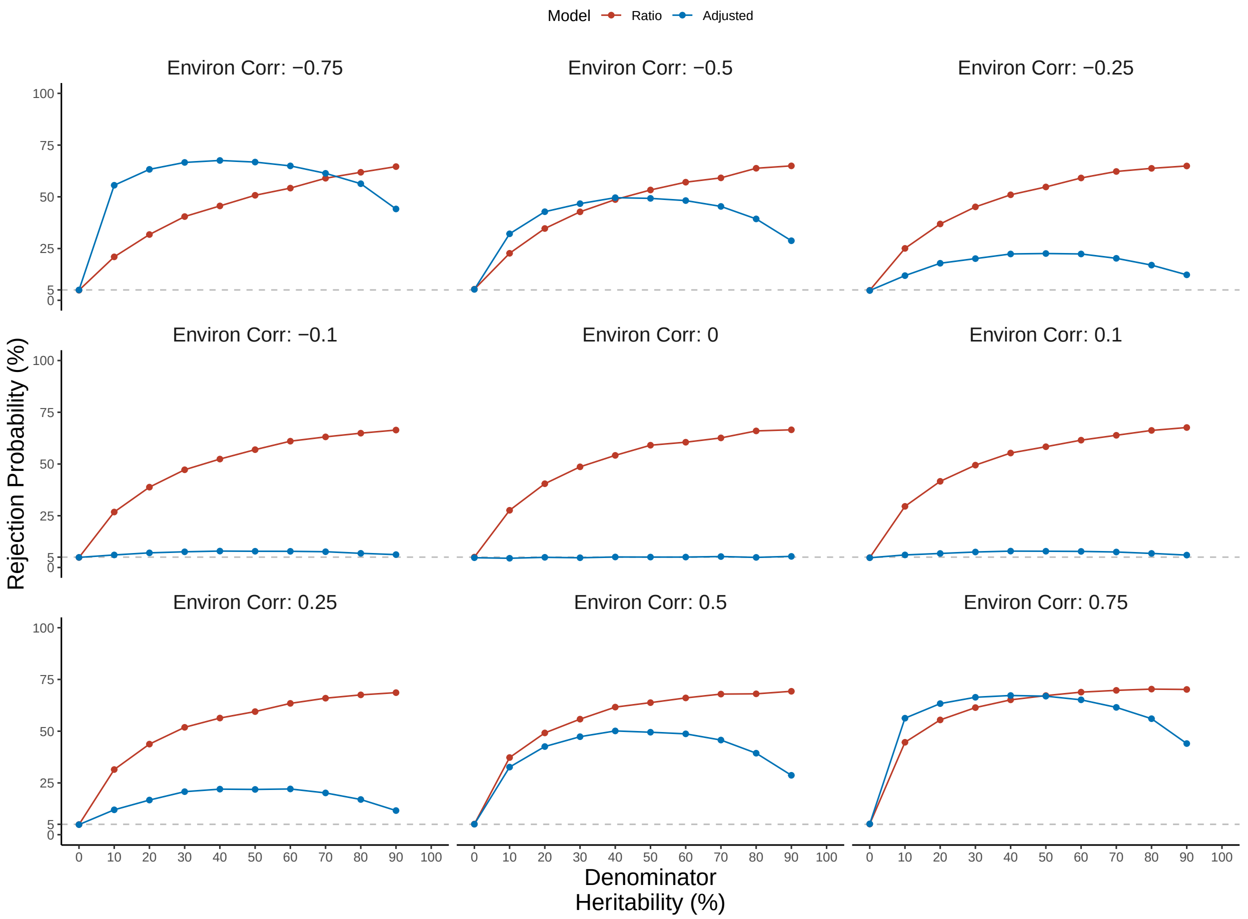

Figure S6: **Rejection probability of the ratio and adjusted models in the presence of environmental collider bias**. Numerator and denominator traits representing weight and height$^2$ were simulated for $N = 10^4$ subjects. Genotype had no direct effect on the numerator, while the denominator had heritability $h^2$, as specified along the X-axis. Environmental correlation between the numerator and denominator traits was set to $\rho$, as specified in the panel heading. Plotted is the rejection probability of the ratio $Y/X \sim G$ and adjusted $Y \sim X + G$ models. Each point is the mean across $R = 500$K simulation replicates.

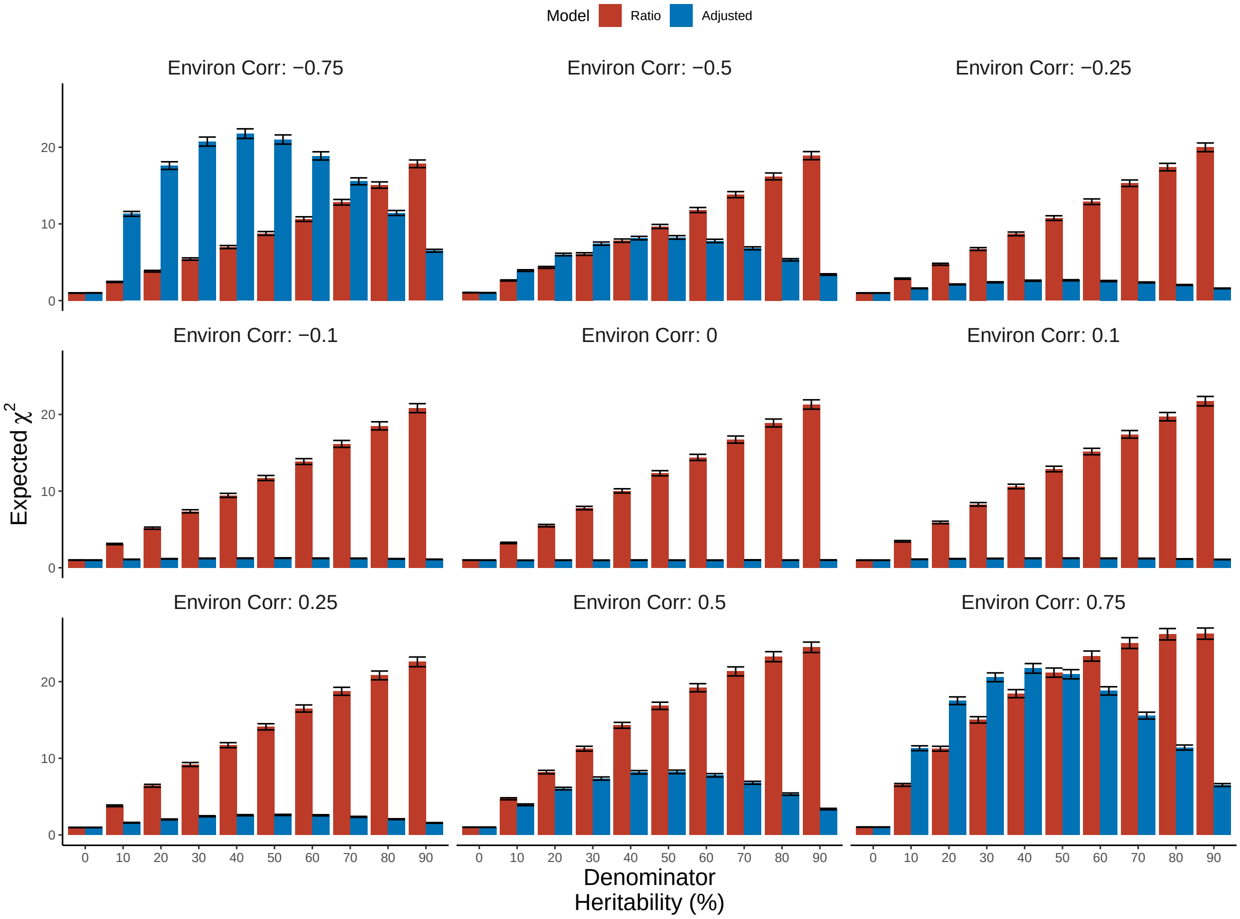

Figure S7: **Strength of association of the ratio and adjusted models in the presence of environmental collider bias**. Numerator and denominator traits representing weight and height$^2$ were simulated for $n = 10^4$ subjects. Genotype had no direct effect on the numerator, while the denominator had heritability $h^2$, as specified along the X-axis. Environmental correlation between the numerator and denominator traits was set to $\rho$, as specified in the panel heading. Plotted is the mean $\chi^2$ statistic, across $R = 500\text{K}$ simulation replicates, of the ratio $Y/X \sim G$ and adjusted $Y \sim X + G$ models. Error bars are 95% confidence intervals.

#### 2.5 Pleiotropy simulation

To investigate the impact of pleiotropy on the operating characteristics of the ratio and adjusted models, numerator and denominator traits were simulated for $n = 10^4$ subjects from the following horizontal pleiotropy model:

$$
\begin{aligned}
Y &= \mu_Y + \sigma_Y\big(G\beta_G + \varepsilon_Y\big), \\
X &= \mu_X + \sigma_X\big(G\alpha_G + \varepsilon_X\big),
\end{aligned}
$$

where $(\mu_Y, \sigma_Y)$ are constants set to match the mean and SD of $Y$ to the empirical mean and SD of weight; $(\mu_X, \sigma_X)$ are constants set to match the mean and SD of $X$ to those of height$^2$; and $G$ represents genotype at $J = 10^2$ variants in linkage equilibrium. Effect sizes were simulated from an infinitesimal model, such that the heritabilities of both $X$ and $Y$ were $h^2$, and the correlation of effect sizes was $\rho$:

$$
\begin{pmatrix} \beta_G \\ \alpha_G \end{pmatrix} \sim N\left\{ \begin{pmatrix} 0 \\ 0 \end{pmatrix}, \begin{pmatrix} h^2/J & \rho \\ \rho & h^2/J \end{pmatrix} \right\}.
$$

Uncorrelated residuals were simulated from a bivariate normal distribution:

$$
\begin{pmatrix} \varepsilon_Y \\ \varepsilon_X \end{pmatrix} \sim N\left\{ \begin{pmatrix} 0 \\ 0 \end{pmatrix}, \begin{pmatrix} 1 - h^2 & 0 \\ 0 & 1 - h^2 \end{pmatrix} \right\}.
$$

Note that under this data generating process, $G$ always has a direct effect on $Y$, thus all rejections are unambiguously true positives.

**Figures S8 and S9** present the power and mean $\chi^2$ statistics of the ratio and adjusted models. In the absence of effect size correlation $\rho = 0$, the ratio and adjusted models have equivalent power. In the presence of positive effect size correlation $\rho > 0$, the adjusted model has a slight but inconsistent power advantage. Meanwhile, in the presence of negative effect size correlation $\rho < 0$, the ratio model has a consistent power advantage, which is most pronounced for $\rho \leq -0.5$. This finding is again consistent with the observations in

Supplementary Section 1.5.3, which notes the (log) ratio model has greater power when $\beta_G$ and $\alpha_G$ have opposite signs.

Although we have presented a simulation study assessing the power of the ratio and adjusted models, we caution that directly comparing power between the two approaches is difficult to justify in general because the two GWAS target different sets of variants. In particular, the ratio model can detect variants associated with either the numerator or the denominator, while the adjusted model targets variants associated with the numerator while holding the denominator fixed. We recommend selecting an association model principally on the basis of the desired interpretation of the effect sizes, and only secondarily on the basis of power.

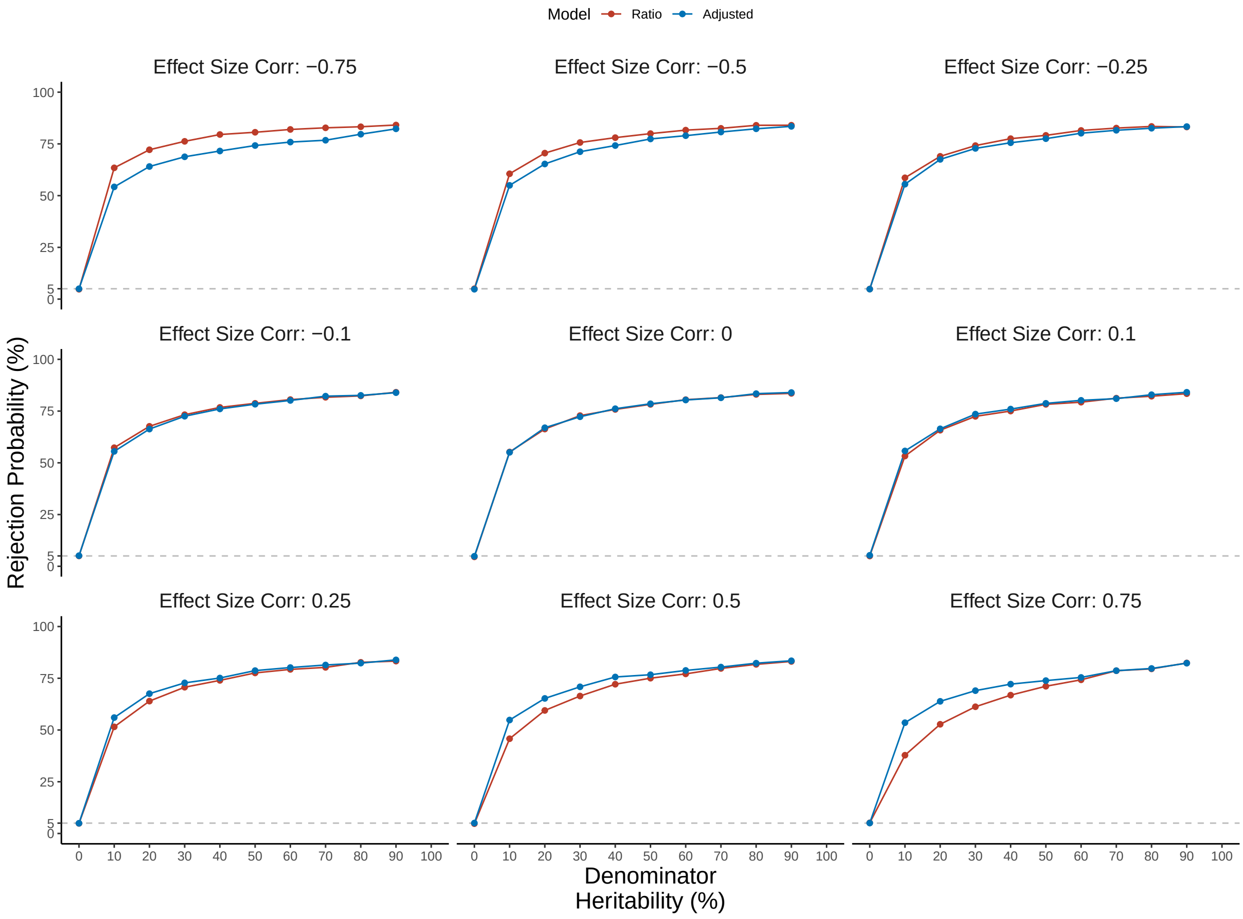

Figure S8: **Power of the ratio and adjusted models in the presence pleiotropy**. Numerator and denominator traits representing weight and height$^2$ were simulated for $N = 10^4$ subjects. Both numerator and denominator had heritable $h^2$, as specified along the X-axis. Effect sizes were simulated from an infinitesimal model, with correlation $\rho$ specified in the panel heading. Plotted is the rejection probability of the ratio $Y/X \sim G$ and adjusted $Y \sim X + G$ models. Each point is the mean across $R = 500$K simulation replicates.

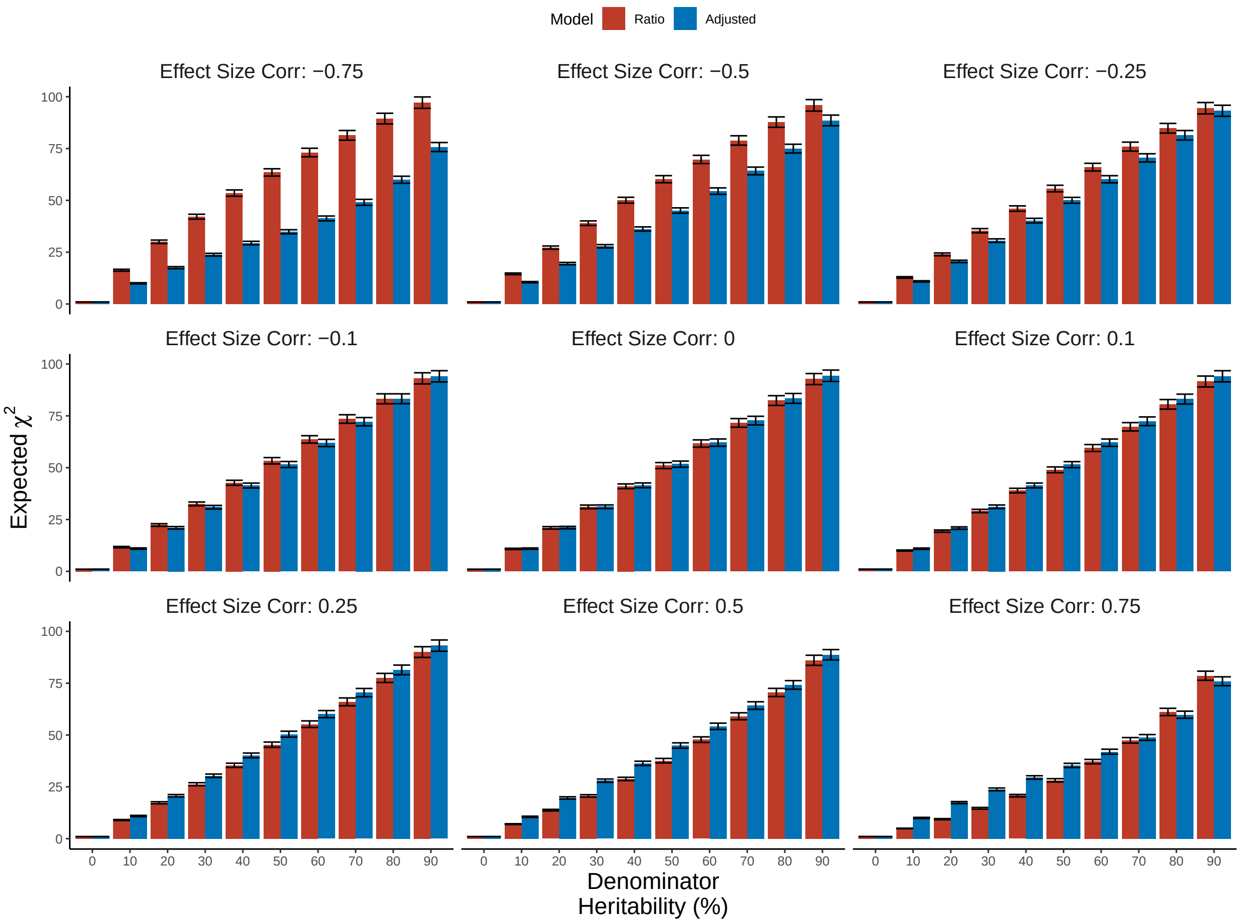

Figure S9: **Strength of association of the ratio and adjusted models in the presence of environmental collider bias**. Numerator and denominator traits representing weight and height$^2$ were simulated for $N = 10^4$ subjects. Both numerator and denominator had heritable $h^2$, as specified along the X-axis. Effect sizes were simulated from an infinitesimal model, with correlation $\rho$ specified in the panel heading. Plotted is the mean $\chi^2$ statistic, across $R = 500$K simulation replicates, of the ratio $Y/X \sim G$ and adjusted $Y \sim X + G$ models. Error bars are 95% confidence intervals.

#### 2.6 Survey of ratio traits

(a) **Ratio Overlap Adjusted**

| Numerator | Denominator | N | Ratio | Ratio-only | Ratio ∩ Adjusted |
|---|---|---|---|---|---|
| Albumin (blood) | Creatinine (blood) | 311122 | 944 | 727 | 217 (22%) |
| Albumin (urine) | Creatinine (urine) | 105501 | 6 | 6 | 0 (0%) |
| Weight | Height | 355627 | 844 | 89 | 755 (89%) |
| FEV1 | FVC | 325220 | 538 | 86 | 452 (84%) |
| Stroke volume | End diastolic volume | 29171 | 6 | 6 | 0 (0%) |
| Phenylalanine | Tyrosine | 200159 | 251 | 54 | 197 (78%) |
| Platelets | Lymphocytes | 52133 | 2 | 2 | 0 (0%) |
| Seated height | Standing height | 355373 | 440 | 330 | 110 (25%) |
| Trunk fat mass | Whole body fat mass | 349533 | 359 | 132 | 227 (63%) |
| Waist circumference | Hip circumference | 356112 | 465 | 116 | 349 (75%) |
| | Average | 243995 | 385 | 154 | 230 (43%) |

(b) **Adjusted Overlap Ratio**

| Numerator | Denominator | N | Adjusted | Adjusted-only | Adjusted ∩ Ratio |
|---|---|---|---|---|---|
| Albumin (blood) | Creatinine (blood) | 311122 | 556 | 314 | 242 (43%) |
| Albumin (urine) | Creatinine (urine) | 105501 | 4 | 4 | 0 (0%) |
| Weight | Height | 355627 | 830 | 70 | 760 (91%) |
| FEV1 | FVC | 325220 | 474 | 49 | 425 (89%) |
| Stroke volume | End diastolic volume | 29171 | 0 | 0 | 0 (.) |
| Phenylalanine | Tyrosine | 200159 | 174 | 10 | 164 (94%) |
| Platelets | Lymphocytes | 52133 | 0 | 0 | 0 (.) |
| Seated height | Standing height | 355373 | 115 | 39 | 76 (66%) |
| Trunk fat mass | Whole body fat mass | 349533 | 240 | 46 | 194 (80%) |
| Waist circumference | Hip circumference | 356112 | 529 | 115 | 414 (78%) |
| | Average | 243995 | 292 | 64 | 227 (68%) |

Table S1: **Overlap of loci detected by the ratio and adjusted models.** Numbers of independent genome-wide significant loci detected by the ratio and adjusted models, decomposed into the count detected exclusively by one model, and the count detected by both models. Since a single locus detected by the ratio model can overlap multiple loci detected by the adjusted model, and vice versa, the overlap analysis is shown two ways. (a) partitions the loci detected by the ratio model into either ratio-only or both, while (b) partitions the loci detected by the adjusted model into either adjusted-only or both. Loci were obtained by clumping full-genome summary statistics at $R^2 \leq 0.10$ and $P \leq 5 \times 10^{-8}$. A locus detected by the ratio model was considered tagged by the adjusted model if within 250kb of an adjusted locus, and vice versa.

| Numerator | Denominator | Genetic Rho (%) | SE (%) | P |
|---|---|---|---|---|
| Albumin (blood) | Creatinine (blood) | -9.80 | 2.80 | 0.00 |
| Albumin (urine) | Creatinine (urine) | -23.50 | 14.90 | 0.12 |
| Weight | Height | 40.50 | 1.60 | 0.00 |
| FEV1 | FVC | 92.20 | 0.40 | 0.00 |
| Stroke volume | End diastolic volume | 73.80 | 10.50 | 0.00 |
| Phenylalanine | Tyrosine | 49.90 | 6.50 | 0.00 |
| Platelets | Lymphocytes | -34.30 | 31.60 | 0.28 |
| Seated height | Standing height | 90.80 | 0.60 | 0.00 |
| Trunk fat mass | Whole body fat mass | 99.30 | 0.00 | 0.00 |
| Waist circumference | Hip circumference | 88.50 | 0.80 | 0.00 |

Table S2: **Genetic correlations between the numerators and denominators of traits analyzed as ratios.** Genetic correlations were calculated from genome-wide summary statistics using LD score regression.

| Numerator | | | | Denominator | | | |
|---|---|---|---|---|---|---|---|
| Trait | Loci | $h^2$ (%) | SE (%) | Trait | Loci | $h^2$ (%) | SE (%) |
| Albumin (blood) | 557 | 11.30 | 0.70 | Creatinine (blood) | 543 | 12.70 | 0.70 |
| Albumin (urine) | 4 | 0.80 | 0.50 | Creatinine (urine) | 10 | 6.00 | 0.60 |
| Weight | 1069 | 23.70 | 0.90 | Height | 4132 | 41.10 | 1.70 |
| $FEV_1$ | 708 | 18.20 | 0.70 | FVC | 851 | 19.00 | 0.80 |
| Stroke volume | 18 | 13.00 | 2.20 | End diastolic volume | 68 | 7.00 | 1.80 |
| Phenylalanine | 79 | 3.60 | 0.70 | Tyrosine | 193 | 7.90 | 1.60 |
| Platelets | 2 | 0.70 | 1.10 | Lymphocytes | 109 | 19.20 | 1.70 |
| Seated height | 1330 | 18.60 | 0.90 | Standing height | 4120 | 41.10 | 1.70 |
| Trunk fat mass | 795 | 21.70 | 0.70 | Whole body fat mass | 789 | 21.80 | 0.70 |
| Waist circumference | 1212 | 19.50 | 0.70 | Hip circumference | 1742 | 19.50 | 0.70 |

Table S3: **Counts of independent genome-wide significant loci and estimated heritability for the numerators and denominators of traits analyzed as ratios.** Loci were obtained by clumping full-genome summary statistics at $R^2 \leq 0.10$ and $P \leq 5 \times 10^{-8}$. Heritability was estimated by LD score regression.

#### 2.7 Effect size correlations

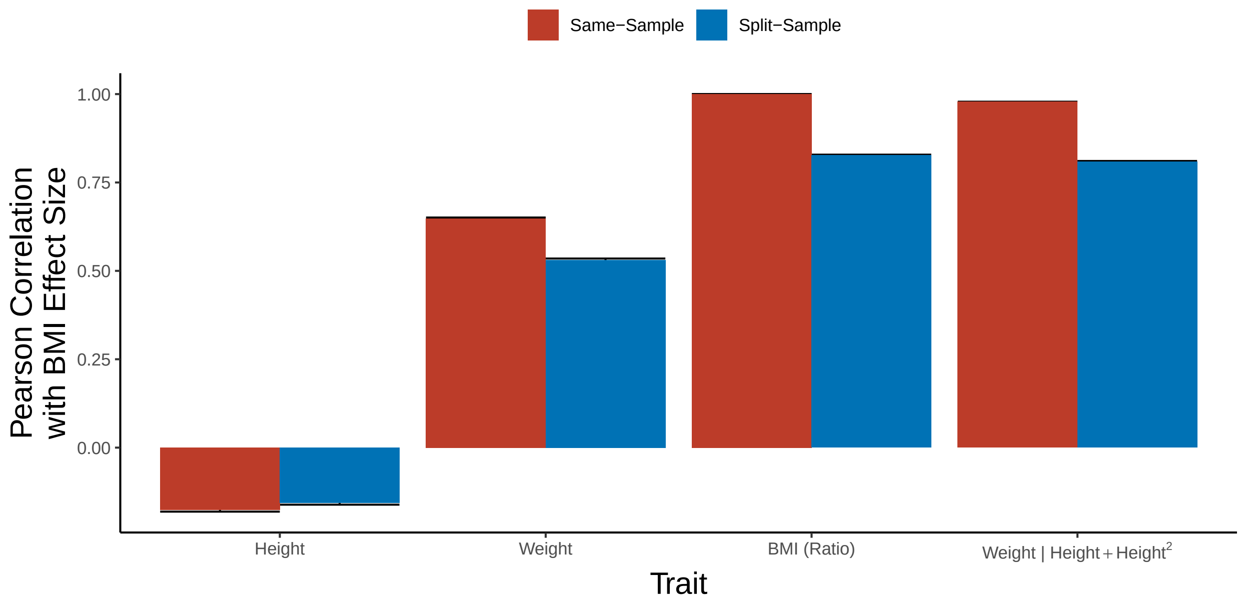

Figure S10: **Correlation of effect sizes with those for BMI, measured either in the same sample or in different samples.** Same-sample refers to correlating effect sizes from two GWAS fit to the same data set, namely the full UKB cohort ($N = 356$K). Split-sample refers to correlating effect sizes estimated in equally-sized independent subsets of the UKB ($N = 178$K each). Effect sizes are correlated across the set of variants significantly ($p \leq 5 \times 10^{-8}$) associated with at least 1 phenotype (BMI, height, weight, weight adjusted for height) in the full sample. Error bars are 95% confidence intervals.

#### 2.8 Biological relevance of ratio and adjusted models

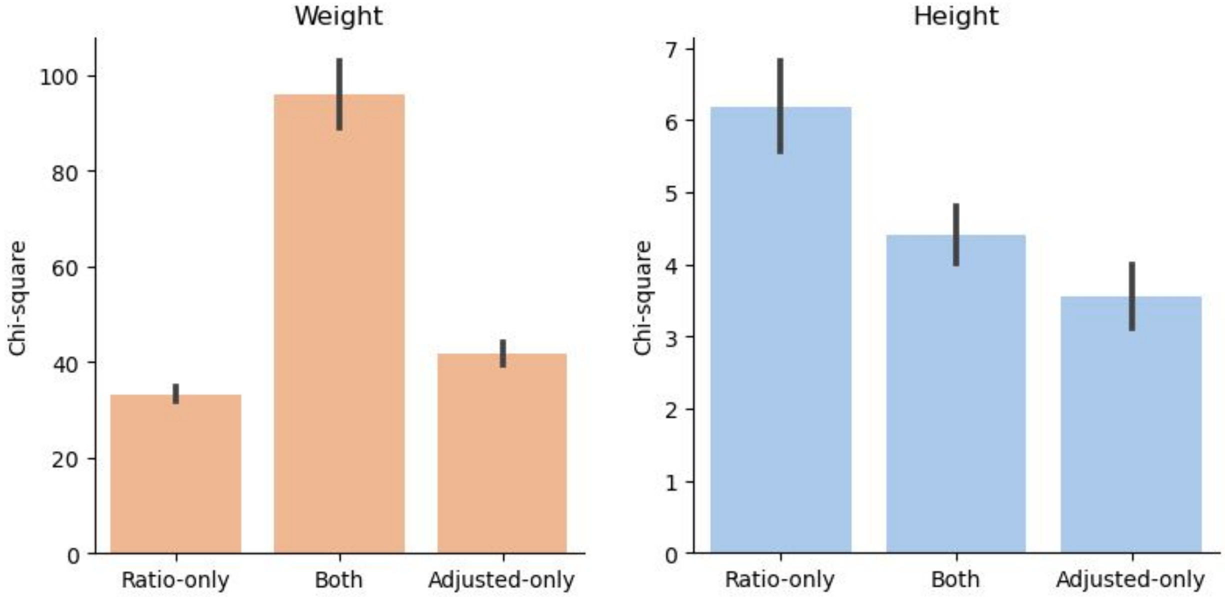

Figure S11: **Adjusted model loci are enriched for numerator signals and ratio model loci are enriched for denominator signals.** GWAS were performed among $N = 356$K unrelated subjects from the UK Biobank. The mean $\chi^2$ statistics for weight (the numerator trait) and height (the denominator trait) partitioned by whether the locus was detected by the ratio model only (i.e. BMI), the adjusted model only (i.e. weight adjusted for height), or both. Error bars are standard errors.

| Trait | $N$ | $h^2$ (SE) | Loci | Adjusted-model average $\chi^2$ (SE) | Ratio-model average $\chi^2$ (SE) | Paired $T$ | $P$ |
|---|---|---|---|---|---|---|---|
| Body fat % [3] | 100,716 | 0.07 (0.01) | 13 | 213.95 (70.31) | 204.59 (68.10) | 2.4169 | 0.0325 |
| Hip circumference [4] | 224,459 | 0.14 (0.01) | 96 | 95.82 (11.82) | 88.59 (11.50) | 8.401 | $4.28 \times 10^{-13}$ |
| T2D [5] | 898,130 | 0.05 (0.002) | 921 | 13.46 (1.52) | 13.41 (1.47) | 0.4371 | 0.6622 |
| CAD [6] | 250,000 | 0.04 (0.003) | 143 | 5.78 (0.92) | 5.79 (0.95) | -0.0927 | 0.9263 |
| cALT [7] | 218,595 | . | 225 | 4.60 (0.60) | 4.63 (0.59) | -0.2675 | 0.7894 |
| LDL cholesterol [8] | 700,000 | 0.09 (0.01) | 4035 | 4.47 (0.14) | 4.39 (0.14) | 2.6782 | 0.0074 |

Table S4: **The adjusted model has more power at loci defined by obesity-related traits.** Loci were obtained by clumping full-genome summary statistics at $R^2 \leq 0.50$ and $P \leq 5 \times 10^{-8}$. Genome-wide summary statistics were not available for chronically-elevated ALT (cALT) [7]. T2D: Type 2 Diabetes, CAD: coronary artery disease.

#### 2.9 Leave-one-chromosome-out polygenic scoring

Following Aschard *et al* [1], data were generated from the 4 data generating processes in **Figure 3(a)**. These cases can be described as:

a. **Complete mediation**, where the effects of genotype $G$ and background $B$ on the phenotype $Y$ are entirely due to $X$.

b. **Non-collider**, where the background $B$ influences both $X$ and $Y$, and $G$ has an effect on phenotype $Y$ but not the covariate $X$.

c. **Collider without direct effect**, where $B$ influences both $X$ and $Y$, and $G$ has an effect on the covariate $X$ but not the phenotype $Y$.

d. **Collider with direct effect**, where $B$ and $G$ both influence $X$ and $Y$.

For the simulation study, $G$ represents genotype at the variant of interest and $B$ represents the cumulative effect of variants **not** in LD with $G$ (i.e. the genetic background). Coefficients were chosen such that the total variation in each variable is 1.0, and the correlation between between $X$ and $Y$ was $\rho \in \{0.00, 0.25, 0.50, 0.75, 0.90\}$. Wherever an edge from $G$ is present, $G$ explains 1% of the variation in the downstream variable (i.e. $h^2 = 1\%$). The simulation was repeated $10^4$ times at each level of $\rho$.

Five estimators are compared:

i. **Adjusted GWAS** of $Y$ on $G$ adjusting for $X$. This estimator is expected to exhibit bias for data generating processes where $X$ is a collider.

ii. **Ratio GWAS** of $Y/X$ on $G$. This estimator is expected to regularly exhibit bias because the association model is misspecified. In practice, the fact that genotype $G$ is rarely expected to act directly on the ratio, as opposed to the individual phenotypes, should be considered before performing a ratio GWAS.

iii. **Conditional GWAS** of $Y$ on $G$ adjusting for $X$ and $B$. This estimator is unbiased and theoretically optimal but requires knowledge of $B$.

iv. **LOCO-PGS GWAS** of $Y$ on $G$ adjusting for $X$ and a LOCO-PGS for $X$. The weights of the PGS were estimated by regressing $Y$ on $B$ within the same sample, then constructing a score $S$ for each subject. Note that in the real data analysis, the PGS weights were estimated in an independent sample. Here, we estimate the PGS weights in the same sample to determine what effect, if any, this has on the performance of the estimator. $S$ emulates a LOCO-PGS because $B$ is not in LD with, and in fact is independent from, $G$.

v. **Non-heritable component GWAS** of $Y$ on $G$ adjusting for the residual after $X$ is regressed on the LOCO-PGS from (iv).
